## Supplementary Figures for "Automatic learning mechanisms for flexible human locomotion"

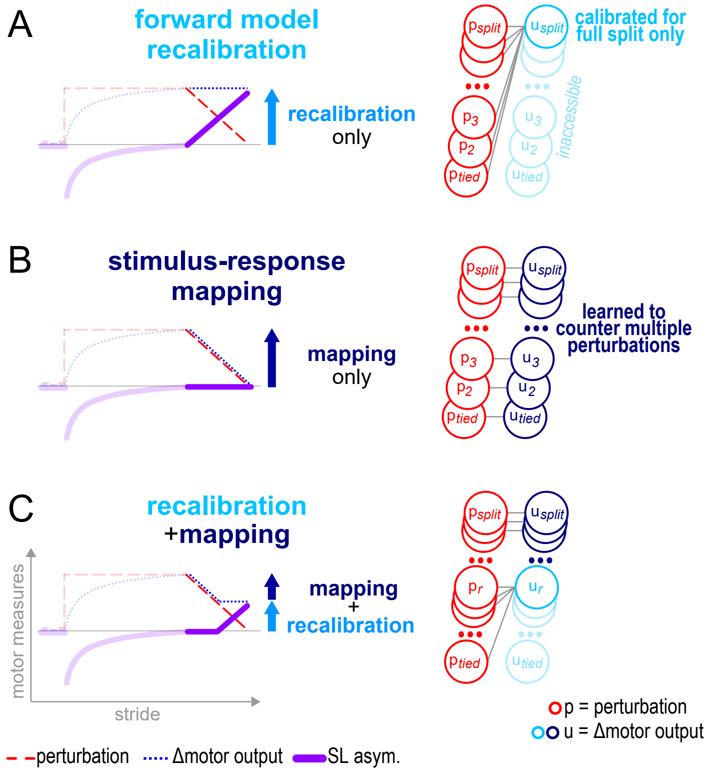


**Figure S1. Experiment 1, conceptual schematic of the learning mechanisms. (A) Forward model recalibration .** By the end of adaptation, the forward model is recalibrated to account for the full split perturbation. The recalibration mechanism can only access the last learnt Δmotor output and uses it regardless of the perturbation (The Δmotor output $u_{split}$, light blue, is used for all perturbations $p_{tied}$, $p_2$, $p_3$, ..., $p_{split}$, red), leading to step length asymmetry aftereffects throughout the Ramp Down. **(B) Stimulus-response mapping.** Mapping can access all Δmotor outputs learnt in adaptation and can select the one matching the perturbation (The appropriate Δmotor output $u_{tied}$, $u_2$, $u_3$, ..., or $u_{split}$ , dark blue, is used in response to perturbations $p_{tied}$, $p_2$, $p_3$, ..., or $p_{split}$, red), leading to no step length asymmetry aftereffect. **(C) Recalibration + mapping.** The Δmotor output can match perturbations accounted for by mapping (The appropriate Δmotor output in the range of $u_r$, $u_{r+1}$, ..., $u_{split}$, dark blue, is used in response to perturbations in the range $p_r$, $p_{r+1}$, ..., $p_{split}$. ), such that step length asymmetry remains zero in the first part of the Ramp Down. For smaller perturbations, the Δmotor output is fixed to the last learnt calibration (The Δmotor output $u_r$, light blue, is used for all smaller perturbations $p_{tied}$, ..., $p_r$), such that step length asymmetry aftereffects begin to emerge. Note that these are simplified illustrations of concepts and not realistic predictions.

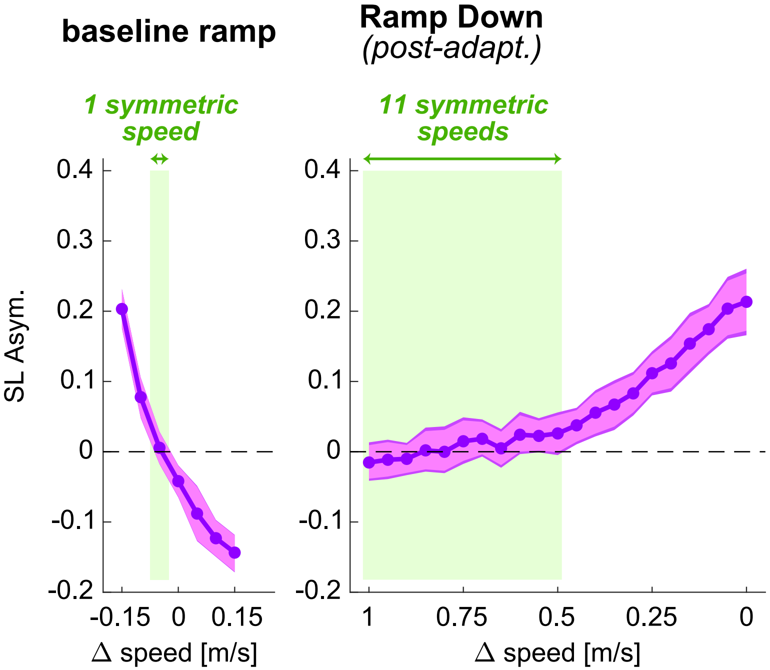


**Figure S2. Experiment 1, step length asymmetry in ramp tasks.** Step length asymmetry (pre-averaged within participant for strides taken at the same speed) as a function of speed (mean, purple curve and dots; 95% CI, light shade; and CI corrected for multiple comparisons, darker shade). Green shaded area represents speeds for which step length asymmetry is not significantly different than zero.

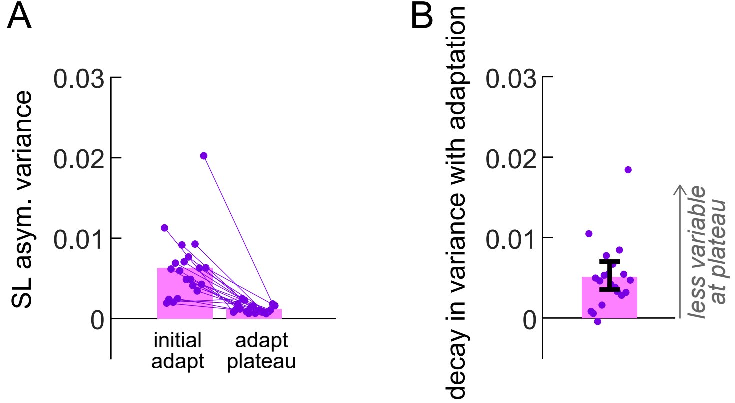


**Figure S3. Experiment 1, variability in adaptation. (A)** Within-participant variance in step length asymmetry in the first and last 30 strides of adaptation. **(B)** Decay in variance between these time points. Bars and error bars: group mean ± CI, circles: individual participants.

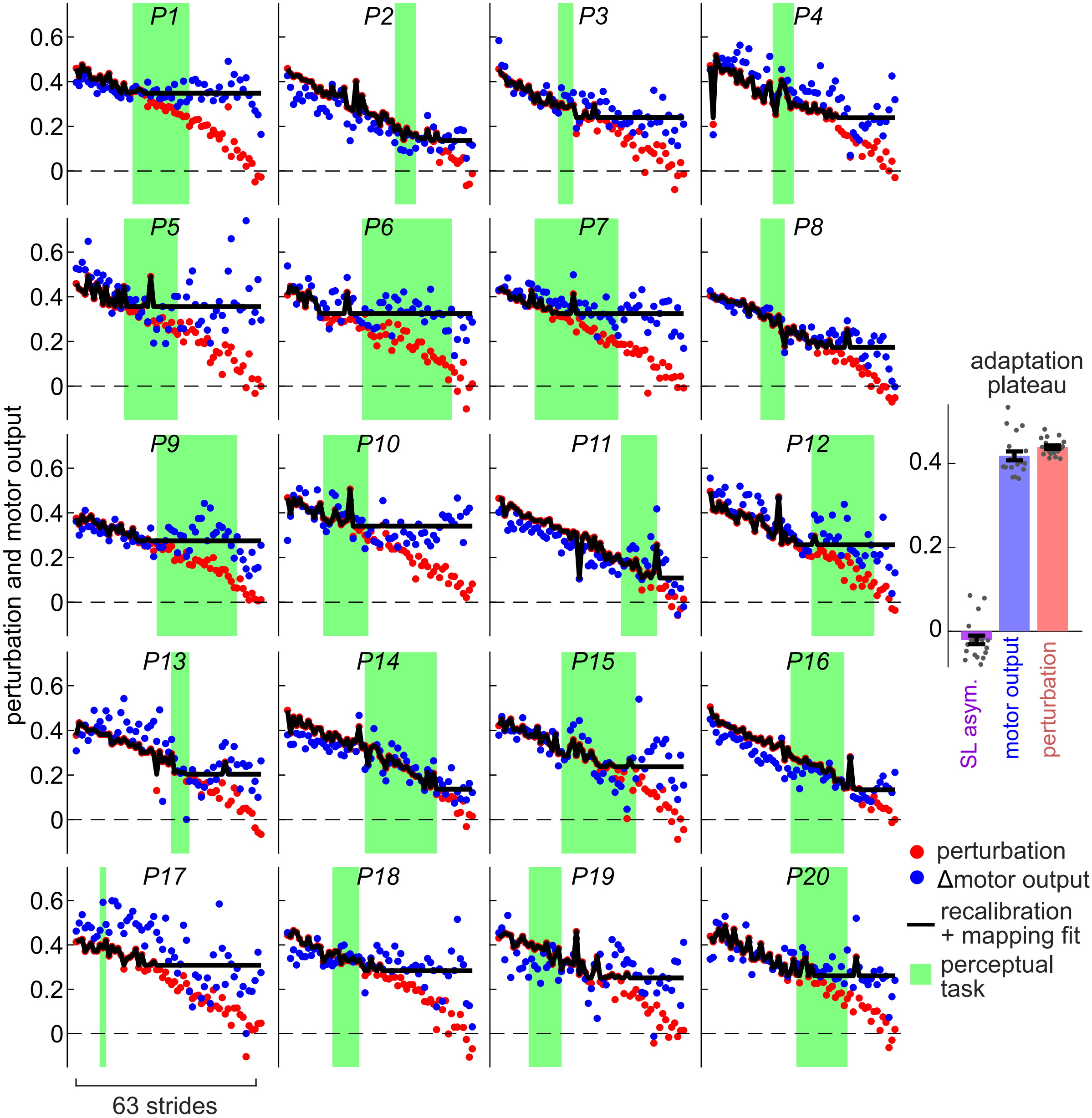


**Figure S4. Experiment 1, individual participants' recalibration + mapping fits and perceptual results.** Perturbation (red circles), Δmotor output (blue circles) and recalibration + mapping model fitted to the Δmotor output (black line), for the Ramp Down task. Green shaded area represents strides between button presses of the perceptual task. The insets show step length asymmetry, Δmotor output, and perturbation at adaptation plateau (mean of the last 30 strides; circles are individual participants and error bars depict group mean ± SE).

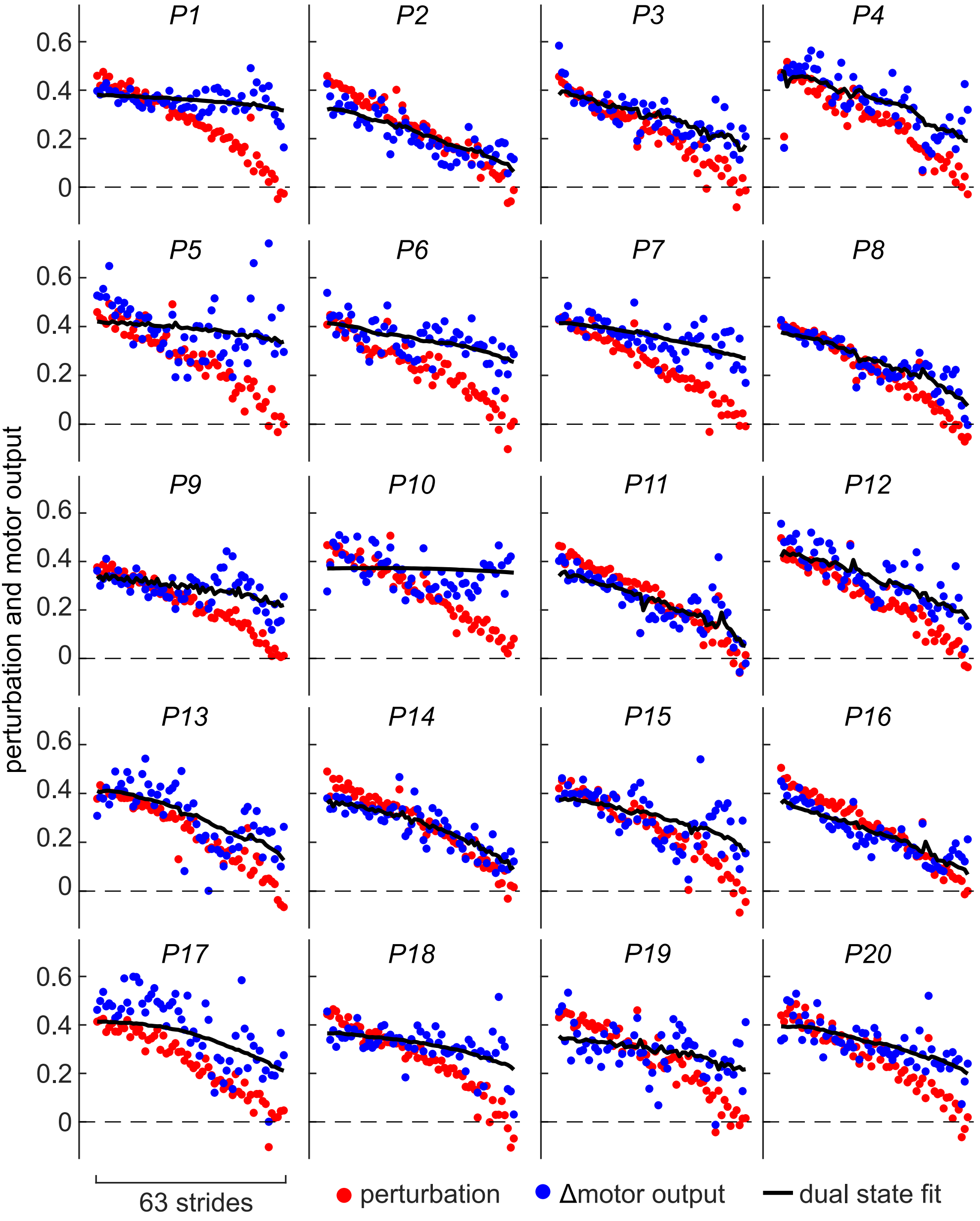


**Figure S5. Experiment 1, individual participants' dual state fits.** Perturbation (red circles), Δmotor output (blue circles) and dual state model fitted to the Δmotor output (black line), for the Ramp Down task.

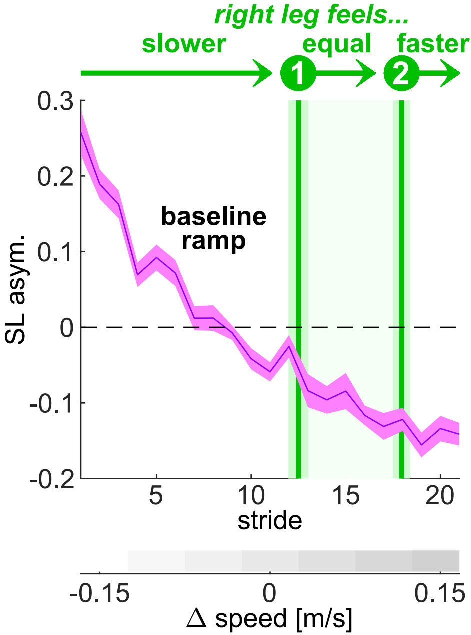


**Figure S6. Experiment 1, baseline ramp perceptual results** (button presses mean ± SE, green vertical lines), overlayed on step length asymmetry (purple line and shade, mean ± SE).

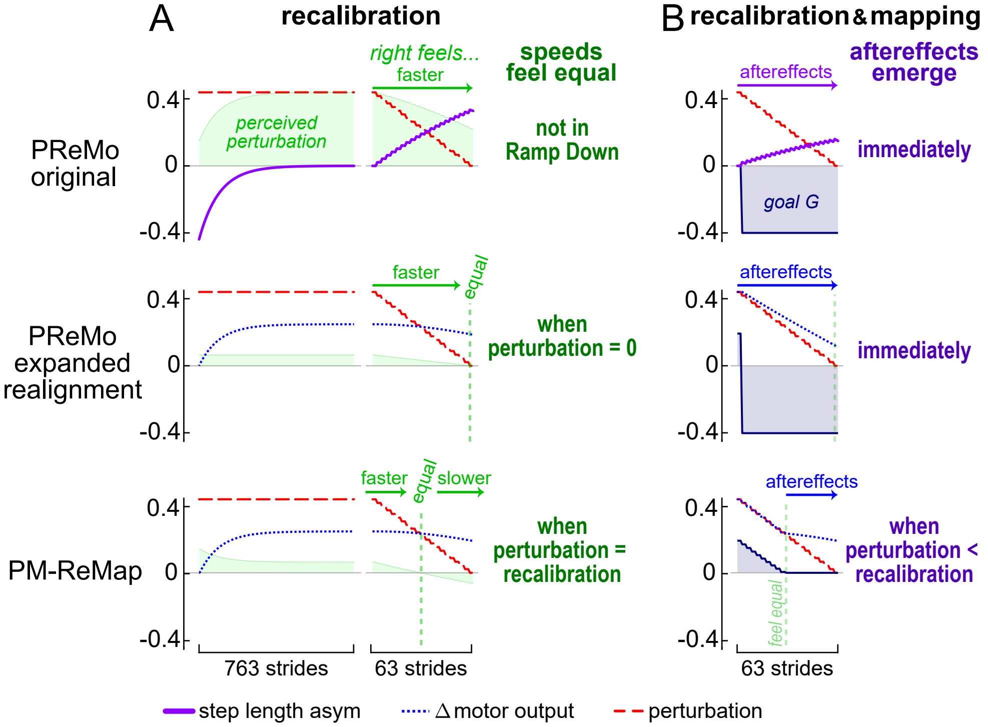


**Figure S7. Experiment 1, perceptual models simulations.** Top: original proprioceptive re-alignment model (PReMo). Middle: expanded version of PReMo that accounts for perceptual realignment. Bottom: perceptuomotor recalibration + mapping model (PM-ReMap). **(A)** Simulations for adaptation by recalibration only. Only PM-ReMap can account for perceptual realignment (belts feel equal halfway through the Ramp Down). **(B)** Simulations for adaptation by both recalibration and mapping. Only PM-ReMap can account for the pattern of motor aftereffects (emerging halfway through the Ramp Down).

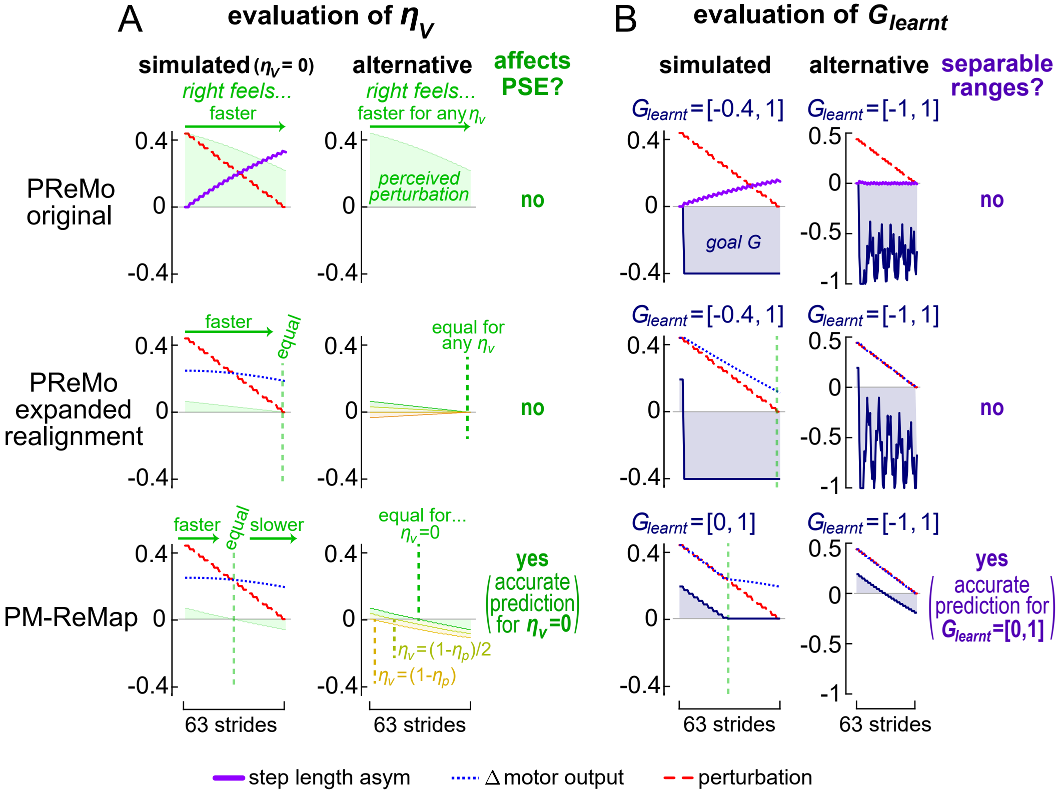


**Figure S8. Experiment 1, evaluation of $\eta_V$ and Glearnt parameters of perceptual models. (A)** Effect of varying $\eta_V$ on the point of subjective equality. $\eta_V$ only affects PM-ReMap, with accurate predictions for $\eta_V$ = *0.* **(B)** Evaluation of why only PM-ReMap can accurately model mapping in the Ramp Down. Only PM-ReMap predicts separable value ranges for the ideal G* in the first versus second halves of the Ramp Down. The first-half range contains learnt values of Δmotor output, resulting in no aftereffects, while the second-half range contains unleart values, leading to aftereffects.

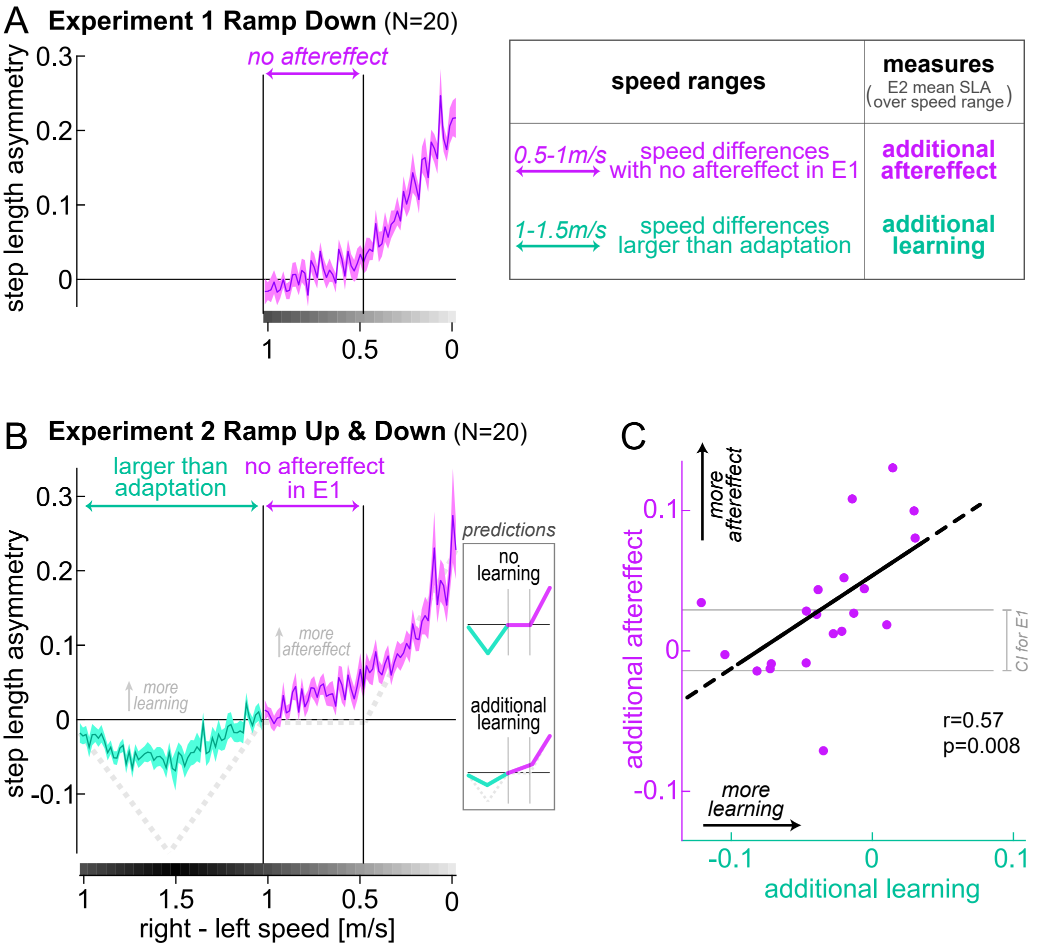


**Figure S9. Experiment 2, evaluation of Ramp Up & Down step length asymmetry differences from Experiment 1. (A) Step length asymmetry time course during the Ramp Down task of Experiment 1** (group mean ± SE). Magenta arrow: range of speeds for which step length asymmetry is not significantly different from zero (no aftereffect). **(B) Step length asymmetry time course during the Ramp Up & Down task of Experiment 2** (entire group mean ± SE). Teal: portion of the task that differs from Exp. 1 because the speed difference is larger than adaptation (>1m/s). Magenta: portion of the task with speeds equal to Exp. 1. The magenta arrow matches that of panel A (speeds with no aftereffect in Exp. 1). Dotted gray line: predicted asymmetry if no additional learning occurred (negative peak magnitude is estimated based on initial adaptation asymmetry). **(C) Experiment 2 individual participants' additional aftereffect versus additional learning** (mean SLA over speed ranges marked by magenta versus teal arrows in panel B). Black line: least square line between the measures. Gray lines: confidence interval for participants in Exp. 1 (CI for the mean SLA over "no aftereffect" range).

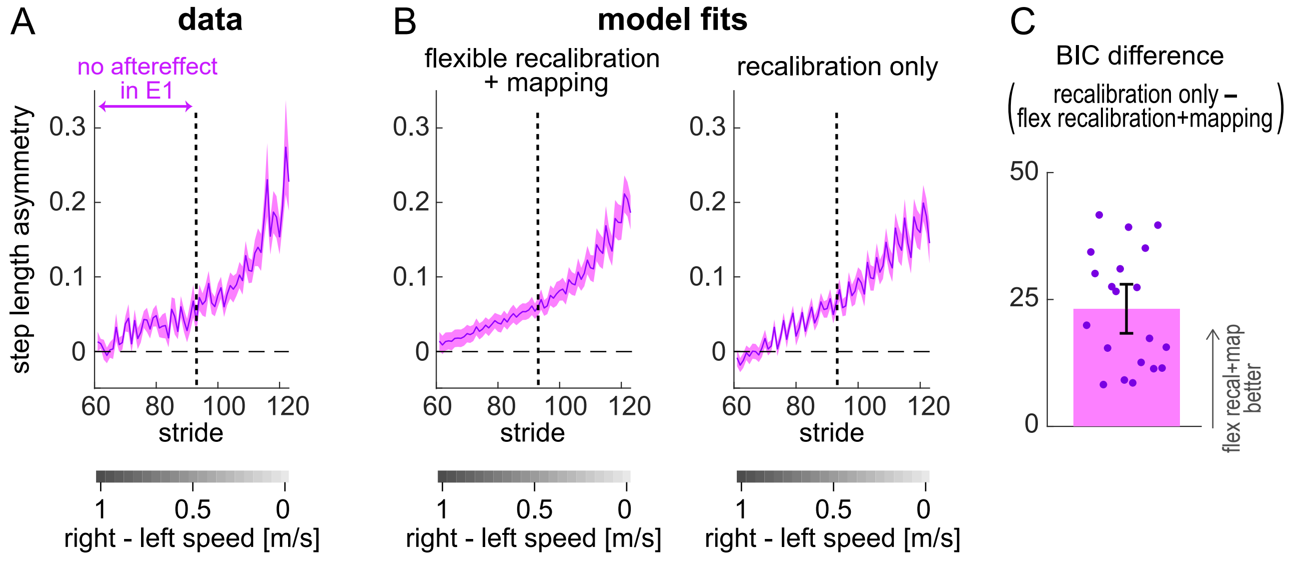


**Figure S10. Experiment 2, second portion of the Ramp Up & Down** (strides 60-12, speed differences 1m/s to 0m/s). **(A)** step length asymmetry data and **(B)** model fits by the flexible recalibration + mapping model (left) and recalibration only model (right). Purple line and shade depict group mean ± SE. Speed configurations to the left of the dashed black vertical line had no aftereffects in E1. **(C)** BIC difference between the recalibration only and flexible recalibration + mapping models. The errorbar depicts group mean and confidence interval, purple dots depict individual participant data. Positve BIC difference indicate that the flexible recalibration + mapping model fits the data better than the recalibration only model.

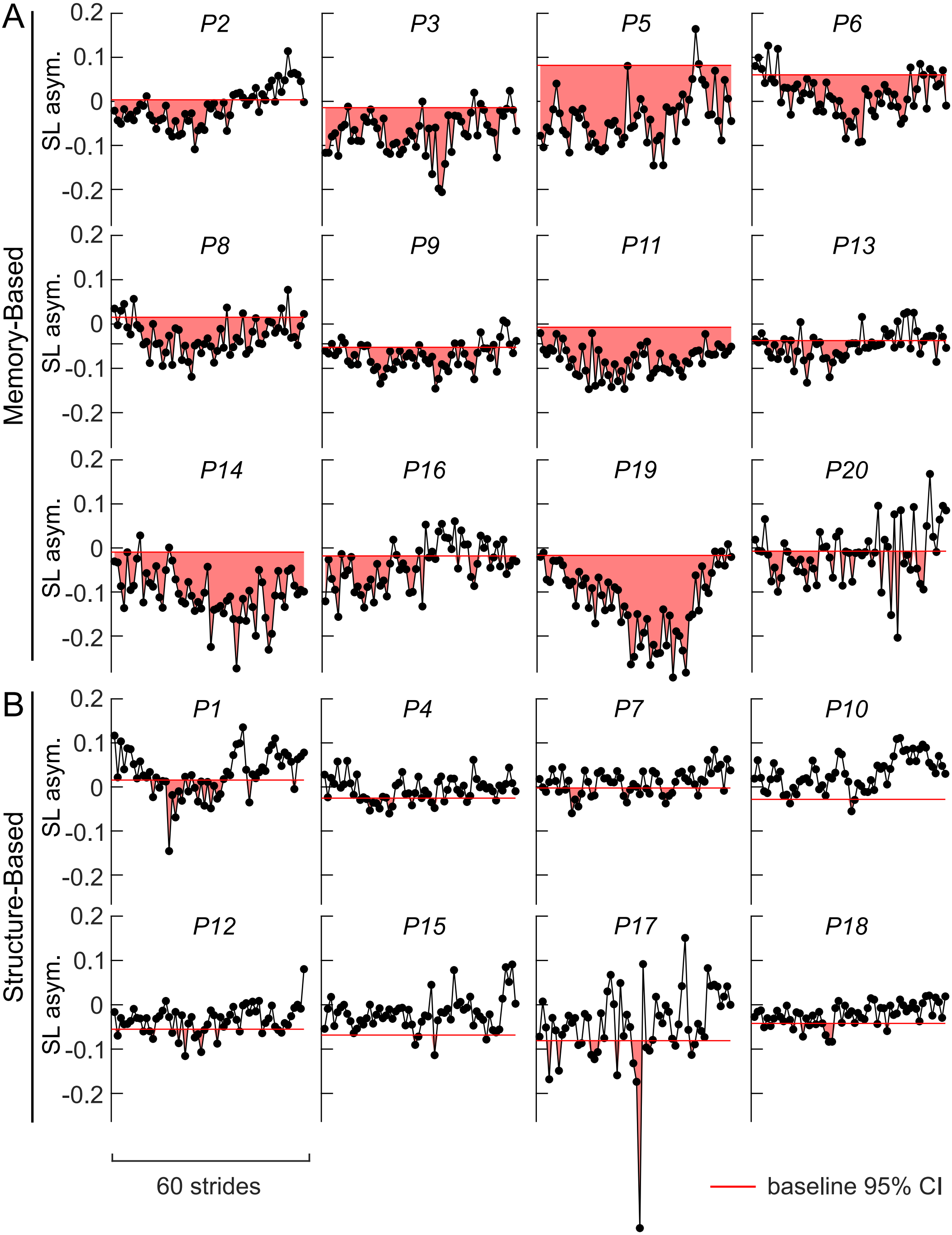


**Figure S11. Experiment 2, individual participants' step length asymmetry in the first portion of the Ramp Up & Down** (speed differences larger than adaptation, teal). The red horizontal line depicts participant's baseline 95% CI (lower bound). Red shaded area represents the difference between task and baseline asymmetry for strides that had a more negative step length asymmetry than baseline.

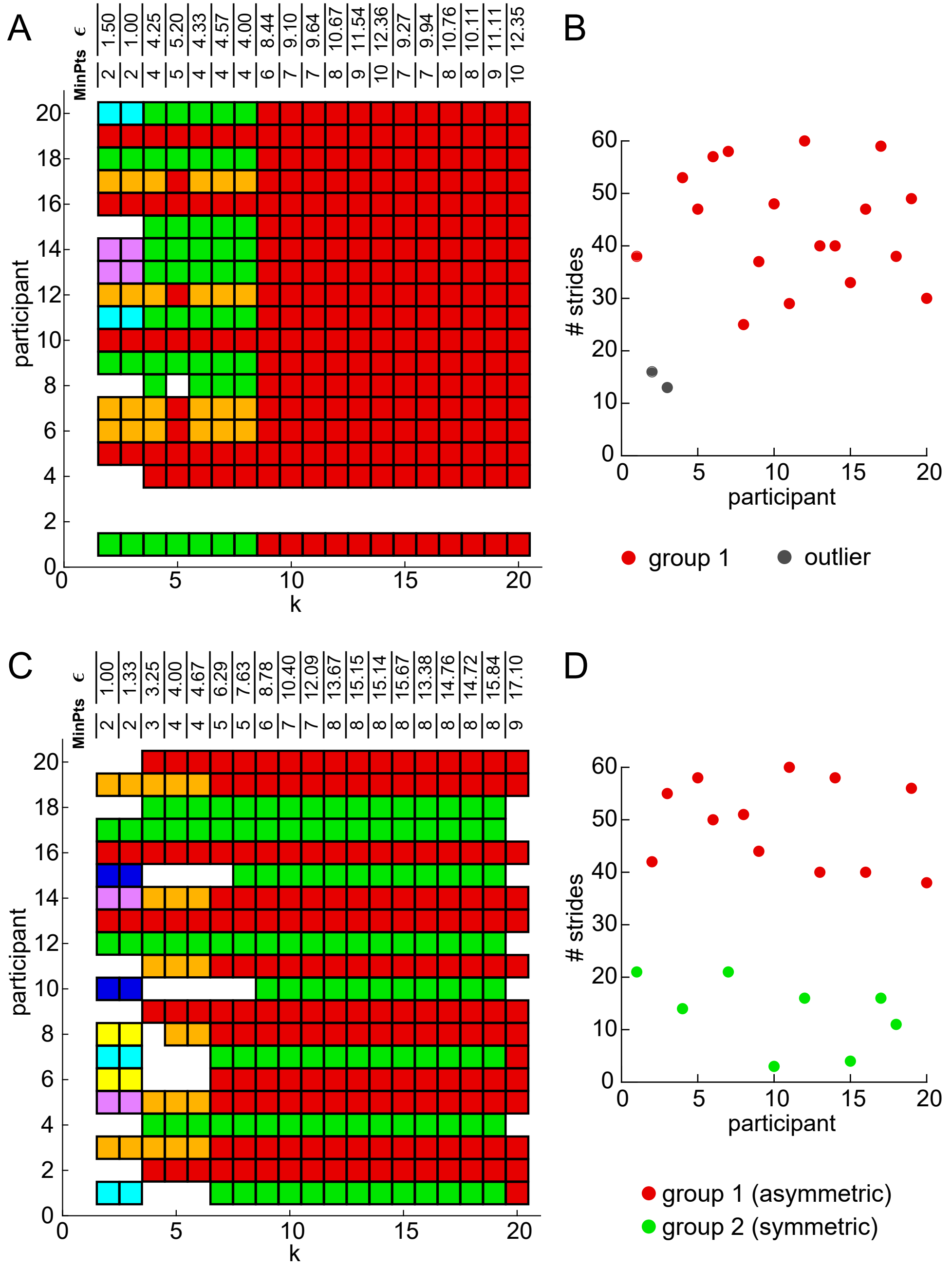


**Figure S12. Primary clustering analysis for Experiment 1 (A&B) and 2 (C&D). (A&C)** Cluster assignment (color of square) for each participant (y axis) computed with each 'k' iteration of the algorithm (x axis). The MinPts and Epsilon parameters computed and used for each 'k' iteration are reported on top of the graph. Different colors represent different clusters, and white spaces represent outliers. The algorithm selected the final cluster to be that for k=9 for both experiments; of note, results were identical for all 'k' iterations from 9 to 20 (A) or 9 to 19 (C). **(B&D)** Measure used for clustering (y-axis, # strides in ramp below/above baseline, see Methods) for each participant (x-axis), color-coded by cluster assignment.

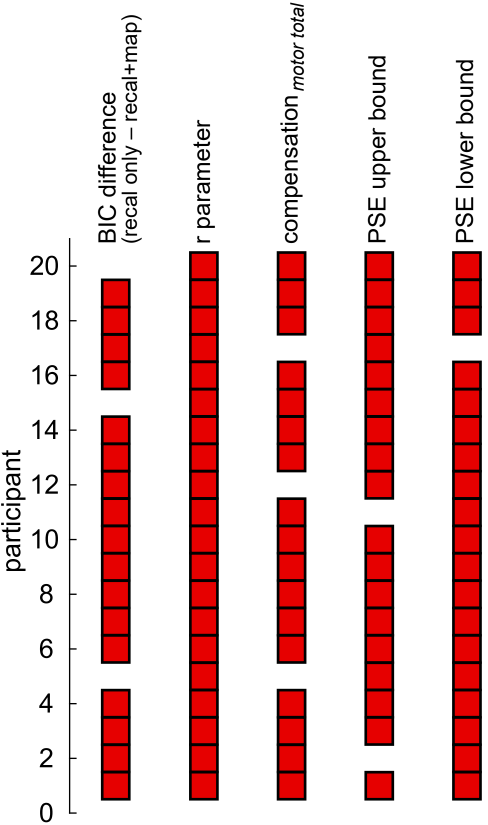


**Figure S13. Secondary clustering analysis for Experiment 1.** Participants clusters (after selection of the best k iteration) computed using different measures (reported above graph). Red boxes indicate participants assigned to cluster 1 and white spaces indicate outliers. For all measures, only one cluster was detected.

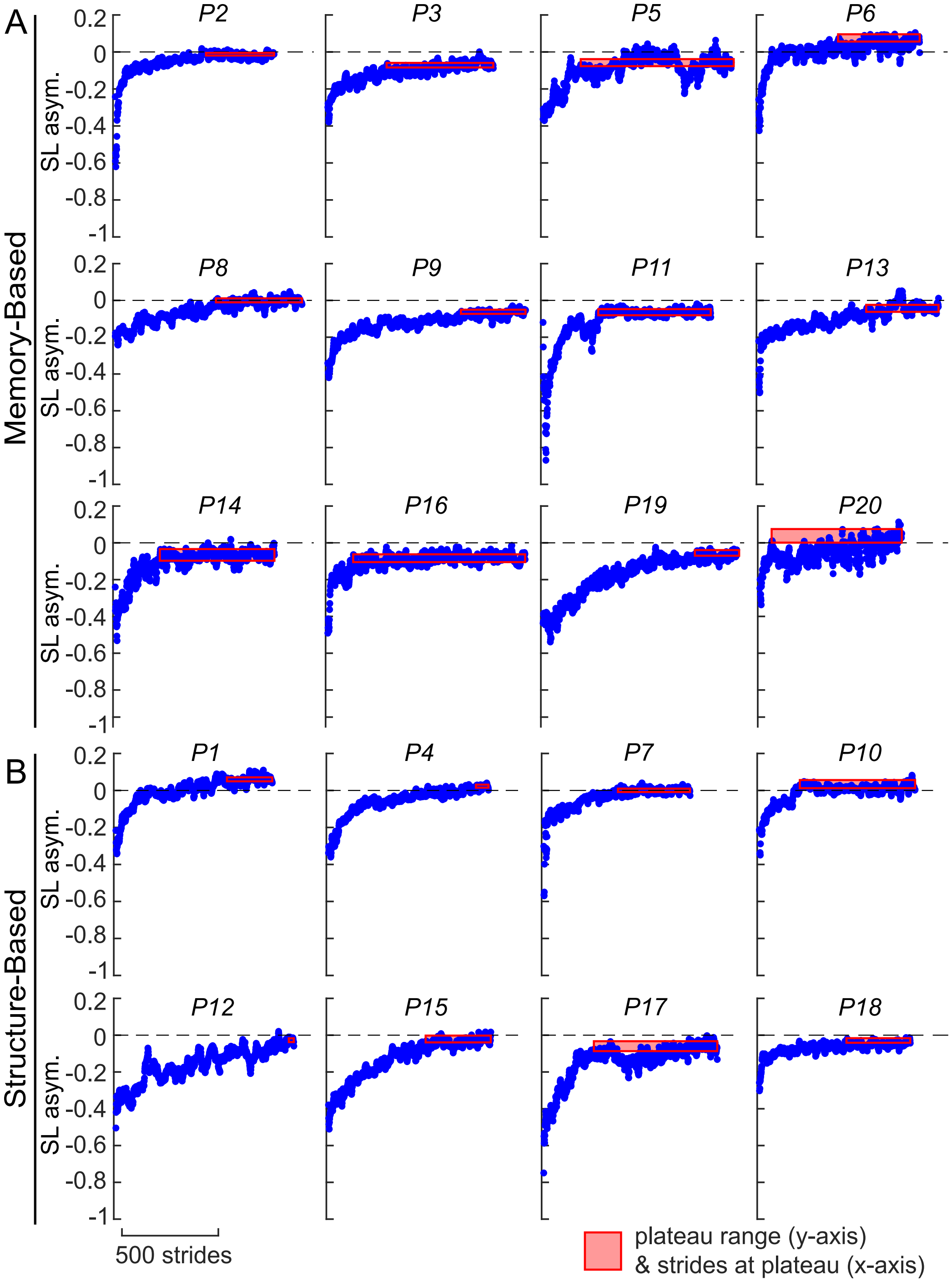


**Figure S14. Experiment 2, individual participants' strides to plateau computation.** Step length asymmetry in adaptation (blue circles), and plateau range (red rectangle; y-axis: plateau mean ± SD, x-axis: strides at plateau).

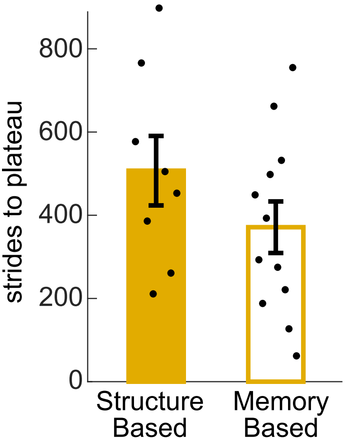


**Figure S15. Experiment 2, comparison between subgroups of strides to plateau measure.** Individual (black circles) and group mean ± SE (bars and error bars) strides to plateau measure divided by subgroup.

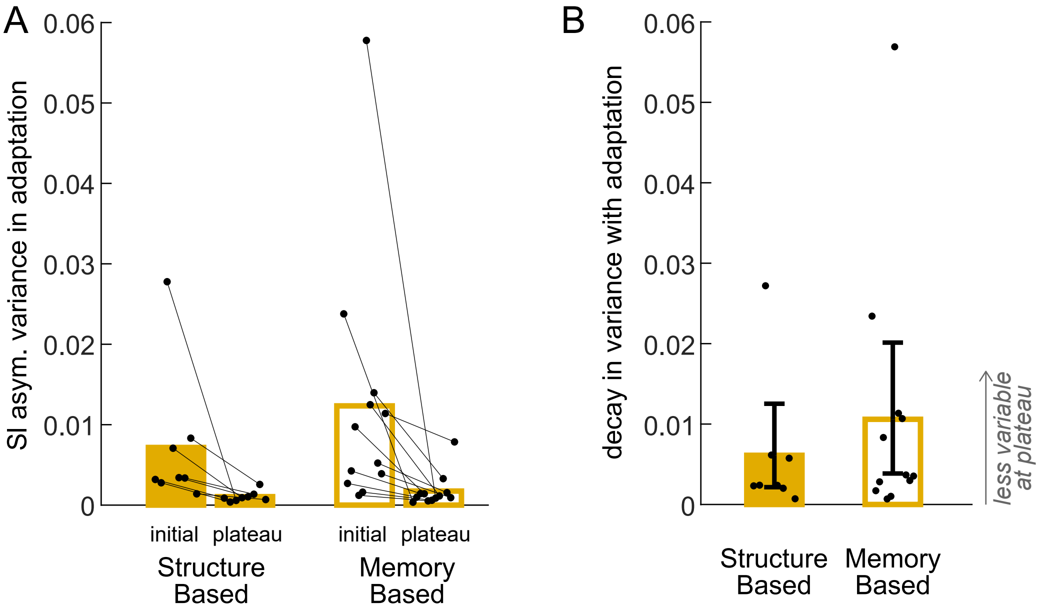


**Figure S16. Experiment 2, variability in adaptation. (A)** Within-participant variance in step length asymmetry in the first and last 30 strides of adaptation, for participants in each subgroup. **(B)** Decay in variance between these time points. Bars and error bars: subgroup mean ± CI, circles: individual participants.
