## Supplementary Appendix for "Automatic learning mechanisms for flexible human locomotion"

[a] Department of Neuroscience, The Johns Hopkins University School of Medicine, Baltimore, MD, 21205, USA; [b] Center for Movement Studies, Kennedy Krieger Institute, Baltimore, MD, 21205, USA; [c] Division of Biokinesiology and Physical Therapy, University of Southern California, Los Angeles, CA, 90033, USA; [d] Neuroscience Graduate Program, University of Southern California, Los Angeles, CA, 90007, USA; [e] Department of Physical Medicine and Rehabilitation, The Johns Hopkins University School of Medicine, Baltimore, MD, 21205, USA

*Corresponding author: Amy J. Bastian.

**This PDF file includes:**

Supporting text:

Ramp Down design (p. 2)

Evaluation and development of perceptual models (p. 5)

Ramp Down comparison between Experiments 1 and 2 (p. 10)

Clustering analysis (p. 17)

Pseudocode for the clustering analysis (p. 19)

Tables S1 to S12 (p. 20)

SI References (p. 32)

Figures S1 to S16 (p. 34)

**Supporting Text**

<u>Ramp Down design</u>

A critical feature of our experimental design is the rate of the Ramp Down following the adaptation phase. The specific design was selected to achieve three main objectives:

1) Minimizing unwanted unlearning or washout by limiting the task duration, because a task that is too slow may eliminate aftereffects regardless of the underlying mechanism

2) Achieving sufficient resolution, with enough measurements of aftereffects across different speeds

3) Ensuring robust measurements at each speed, collecting sufficient strides per condition for reliable estimates

To achieve these goals, we evaluated previous literature. However, to our knowledge, existing paradigms with gradual perturbation ramp-downs post-adaptation do not fully align with these objectives. Most paradigms were designed to minimize aftereffect errors or participants' awareness of perturbations and employed slow ramp-downs (e.g., 10-minute protocols in earlier walking adaptation work) (Herzfeld et al., 2014; Orban De Xivry and Lefèvre, 2015; Roemmich and Bastian, 2015; Schlerf et al., 2013). Thus, we relied on alternative studies to inform specific aspects of our design.

We used the work by Leech et al. to decide the specific speeds to test in our Ramp Down design (Leech et al., 2018a). As described in Box 2, their "speed match" task provided preliminary evidence for flexible mapping mechanisms and informed our hypotheses and design in multiple ways. To determine an appropriate resolution for the Ramp Down speeds, we analyzed their baseline perceptual results to assess how small a difference in belt speeds participants could perceive. Within participants, the belt speed differences perceived as "equal speed" varied across the three baseline iterations of the speed-match task, spanning a range of 0.1 m/s on average. Assuming the true point of subjective equality lies at the midpoint of this range, this indicates that participants can detect belt speed differences of 0.05 m/s or larger. Based on this, we designed the Ramp Down protocol to decrease the speed difference in intervals of 0.05 m/s, as this resolution was sufficient to capture perceptual realignment.

This interval yielded 11 distinct speed conditions, consistent with the range used in various generalization studies (Abeele and Bock, 2001; Leech et al., 2018b; Malfait et al., 2005; Tanaka and Sejnowski, 2015; Taylor and Ivry, 2013). As such, this resolution is sufficient to capture not only perceptual realignment, but also the pattern of motor aftereffects across perturbation sizes. We chose to collect three strides at each speed condition to align with standards established in the split-belt adaptation literature; specifically, three strides are the minimum number consistently used to measure post-adaptation aftereffects, as typically observed in catch trials (Leech et al., 2018a; Rossi et al., 2019; Vazquez et al., 2015). Our Ramp Down design consisted of 63 speed configurations (11 speed conditions × 3 strides each) and lasted 1 minute and 20 seconds on average.

We confirmed that this duration was sufficient to preserve aftereffects based on previous work. Motor and perceptual aftereffects last for several minutes after adaptation to a 3:1 split-belt perturbation (Kambic et al., 2023; Leech et al., 2018a; Vazquez et al., 2015). To obtain specific estimates, we reanalyzed data from Leech et al. and found that both motor and perceptual aftereffects persist for over 4.5 minutes into washout (see *Analysis of aftereffect decay in Leech et al.* below). Of note, we expect less washout in our Ramp Down than in the ascending speed match tasks used by Leech et al. (as discussed, ascending speed match tasks involve perturbations opposite to those learned during adaptation). Therefore, we believe our selected Ramp Down rate was fast enough to minimize forgetting of the aftereffects. Our control experiments replicated the behavior observed in the Ramp Down using speed match tasks lasting 30 seconds, further supporting the robustness of our findings across varying durations.

Finally, as schematized in Figure 1, our Ramp Down design is more robust against misinterpretations than previous approaches that focus solely on aftereffect magnitude. First, it enables the dissociation of the recalibration + mapping hypothesis from the recalibration-only hypothesis, even under conditions of partial washout. Specifically, partial forgetting would reduce the magnitude of aftereffects but would not alter their pattern of emergence (for the recalibration + mapping hypothesis, aftereffects would still emerge halfway through the task, whereas for the recalibration-only hypothesis, they would still appear immediately). Second, if the task duration were too slow and resulted in complete forgetting, aftereffects would be absent throughout the entire ramp. This pattern would be inconsistent with both the recalibration + mapping and recalibration-only hypotheses, serving as a clear indicator that forgetting has confounded the results.

*Analysis of aftereffect decay in Leech et al.*

We reanalyzed data from Leech et al. to evaluate the time course of aftereffect washout (Leech et al., 2018a). We evaluated perceptual aftereffects measured as the baseline-subtracted belt speed difference at the end of each of the six post-adaptation speed match tasks (right – left). We evaluated motor aftereffects measured as step length asymmetry averaged over the five strides following each speed match task. Using a bootstrap analysis equivalent to that described for Experiment 1 (that comparing step length asymmetry in the ramp tasks to zero), we compared these aftereffects to zero to determine when they were last significant. Perceptual aftereffects were last significant in the fifth task, performed 8 minutes into washout (perceptual aftereffect in $5^{th}$ task = 0.035 [0.006 0.061], in $6^{th}$ task = -0.003 [-0.041, 0.037], mean [CI]). Motor aftereffects were last significant following the fourth task but decayed before the fifth task, as confirmed by an additional analysis of the last five strides in that epoch (mean step length asymmetry in first 5 strides after $4^{th}$ task = 0.058 [0.022, 0.091], last 5 strides before the $5^{th}$ task = 0.027 [-0.009, 0.060], first 5 strides after $5^{th}$ task = 0.023 [-0.013, 0.058], mean [CI]). Thus, motor aftereffects decayed between the end of the fourth task and the start of the fifth task – i.e., between 4.5 and 8 minutes into washout. For all statistical analysis, we use False Discovery Rate to correct for multiple comparisons (m=6 for the family of 6 perceptual aftereffect tests, and m=5 for the family of 5 motor aftereffect tests) (Benjamini and Yekutieli, 2005).

**Supporting Text**

### Evaluation and development of perceptual models

*PReMo variables equivalents for walking adaptation*

The first step in evaluating performance of the proprioceptive re-alignment model (PReMo) for walking adaptation was to establish equivalencies between its variables – focused on reaching adaptation – and our walking adaptation variables (Tsay et al., 2022, 2021). While the model was originally developed for reaching adaptation in response to visual-proprioceptive discrepancies, the authors provided an example for how the model can be applied to force-field adaptation. We based our evaluation of PReMo on this example because, like force-field adaptation, split-belt adaptation is driven by a mechanical perturbation. Mechanical perturbations differ from visual perturbations because they introduce a mismatch between predicted and actual sensory outcome of a movement, rather than a mismatch between different sensory modalities.

The PReMo model for force-field adaptation contains the following variables:

- $x_p$= proprioceptive observation of hand position (actual position perturbed by force-field)
- $x_v$ = visual observation of hand position (if visual feedback is absent, $x_v$ is not used and variables that depend on $x_v$ decay back to zero)
- $G$ = sensory prediction for hand position = reaching goal (target plus any aiming strategies)
- $\sigma_p^2, \sigma_v^2, \sigma_u^2$ = proprioceptive, visual, and sensory prediction uncertainties
- $K$ = learning rate

These definitions reveal key principles that we use to apply the model to walking adaptation. First, Tsay et al. clearly defines that $x_v$ is not used if there is no visual feedback, as in our task. Second, for mechanical perturbations, $x_p$ represents the actual outcome of a movement, including the effect of the perturbation on the motor output, so that it corresponds to our step length asymmetry measure. Third, in the absence of explicit strategies, $G$ equals the target value of $x_p$ that eliminates error, corresponding to zero step length asymmetry in our task. As walking adaptation is not thought to involve aiming strategies (Long et al., 2016; Malone and Bastian, 2010; Roemmich et al., 2016) (also see Experiment 2), we hypothesized $G = 0$. However, we also tested whether $G$ may represent the stimulus-response mapping mechanism described in our study.

We apply the PReMo model to walking adaptation using the following variables:

- $x_p$= proprioceptive observation of step length asymmetry = actual step length asymmetry

- $G$ = sensory prediction for step length asymmetry = mapping-related goal
- $\sigma_p^2, \sigma_u^2$ = proprioceptive and sensory prediction uncertainties
- $K$ = learning rate

*Original PReMo simulations*

We used simulations to demonstrate that the original PReMo model cannot capture the patterns of motor and perceptual behaviors observed in the Ramp Down. Note that the equations for perceived and actual step length asymmetry evaluated at adaptation plateau imply no perceptual realignment. This is because adaptation plateaus when the perceptual error is zero, and substituting this condition into the equations above results in no difference between perceived and actual step length asymmetry or perturbation at adaptation plateau:

*Perceived step length asymmetry at plateau:* $x_p^{per}(plateau) = G(plateau)$

*Actual step length asymmetry at plateau:* $x_p(plateau) = G(plateau)$

*Perceived perturbation:* $p^{per}(plateau) = p(plateau)$

This prediction contrasts with our results, demonstrating that the original PReMo model cannot account for the perceptual realignment observed in our data.

We use simulations to substantiate this finding and additionally demonstrate that PReMo fails to account for the motor behavior in the task (Fig. S7A-B, top). We simulate the entire paradigm defining the perturbation $p(k)$ as belt speed difference times the average $p_{plateau}$ across participants, using the average number of strides per epoch. We set $W_p = \frac{\sigma_u^2}{\sigma_u^2 + \sigma_p^2} = \frac{1}{3}$ and $K$=0.03, matching the initial conditions explained in the main text (Zhang et al., 2024). We first simulated adaptation of step length asymmetry via recalibration only ($G = 0$) and confirmed that the model predicts no perceptual realignment (Fig. S7A top, perceived and actual perturbation are equal at the end of adaptation). To model mapping, we evaluated the value of $G$ that would lead to zero step length asymmetry:

*Ideal goal:* $G^* = x_p^{per}(k) - \frac{1}{K}\left(p(k+1) - x(k)\right)$

We simulated the Ramp Down only (see main text), setting $x_p(0) = G(0) = 0$ (reflecting zero asymmetry at adaptation plateau). Setting $G(t+1) = G^*$ for all $t$ led to approximately zero step length asymmetry as predicted (Fig. S8B, top right; small differences are because $G^*$ is computed on previous stride observations).

Capturing the Ramp Down behavior requires setting $G = G^*$ for the first half but not the second, with a biologically plausible rationale for why first-half $G^*$ values are accessible (e.g., in memory) while second-half $G^*$ values are not. However, $G^*$ values overlap fully, ranging from -1.00 to -0.38 in the first half and from -0.97 to -0.41 in the second. This overlap prevents the model from limiting $G$ in a way that replicates our results (Fig. S8B top left shows an example simulation with capped values, leading to aftereffects that emerge immediately albeit growing slowly). In sum, PReMo cannot account for the mapping behavior observed in our data or the perceptual realignment.

*Iterative simulations for the development of PM-ReMap*

We iteratively addressed limitations of the PReMo model using simulations, progressing towards the development of a perceptuomotor recalibration + mapping (PM-ReMap) that can capture our Ramp Down data.

We first addressed the limitation that that PReMo cannot capture perceptual realignment in our study as it assumes it arises from mismatches between sensory modalities, such as vision and proprioception. Using our framework (Rossi et al., 2021), we redefined the error signal driving perceptual realignment to reflect mismatches between the perturbed movement outcome and motor command for mechanical perturbations. Specifically, we reinterpreted the model variables to align with the specific mismatch introduced by the perturbation: $x_p$ represents unperturbed motor output, and $x_v = x_p - p$ represents perturbed movement information (here, step length asymmetry). This adjustment captures perceptual shifts driven by sensory prediction errors without altering the model equations:

*Integrated estimate for motor output*: $$x_p^I(k) = \frac{\sigma_u^2}{\sigma_u^2+\sigma_p^2} x_p(k) + \frac{\sigma_p^2}{\sigma_u^2+\sigma_p^2} G(k)$$

*Integrated estimate for step length asymmetry*: $$x_v^I(k) = \frac{\sigma_u^2}{\sigma_u^2+\sigma_v^2} x_v(k) + \frac{\sigma_v^2}{\sigma_u^2+\sigma_v^2} G(k)$$

*Perceptual shift for motor output:* $$\beta_p(k) = \eta_p \left( x_v^I(k) - x_p^I(k) \right)$$

*Perceptual shift for step length asymmetry:* $$\beta_v(k) = \eta_v \left( x_p^I(k) - x_v^I(k) \right)$$

*Perceived motor output:* $$x_p^{per}(k) = x_p^I(k) + \beta_p(k)$$

*Perceived step length asymmetry:* $x_v^{per}(k) = x_v^I(k) + \beta_v(k)$

*Perception of belt speed difference:* $\Delta^{per}(k) = x_p^{per} - x_v^{per}$

*Implicit adaptation of motor output:* $x_p(k+1) = x_p(k) + K\left(G(k) - x_p^{per}(k)\right)$

We set $W_p = \frac{\sigma_u^2}{\sigma_u^2+\sigma_p^2} = \frac{1}{3}$ and $K = 0.03$ as before, and $W_v = \frac{\sigma_u^2}{\sigma_u^2+\sigma_v^2} = \frac{1}{3}$ to match $W_p$ because both signals are sensed proprioceptively. We set $\eta_p = 0.56$ to align with the average $r$ parameter from the recalibration + mapping model across participants ($\eta_p$ reflects implicit adaptation at plateau when $W_p = W_v$) We tested different values for $\eta_v$ which affects perception but not movement.

The simulation of adaptation via recalibration only ($G = 0$) showed perceptual realignment at adaptation plateau, addressing a limitation of the original model (Fig. S7A, middle row, perceived perturbation, green, is smaller than actual perturbation, red). However, it failed to account for the Ramp Down perceptual results, inaccurately predicting that belt speeds feel equal when they are actually equal (Fig. S7A, middle row, perceived perturbation decays alongside actual perturbation and converge to zero at the end of the Ramp Down). This occurred regardless of the value of parameter $\eta_v$ (Fig. S8A, middle row, value of $\eta_v$ affects the slope of perceived perturbation but not its intercept with the x-axis). This occurs because, under the retained PReMo equations, $\beta_p$ and $\beta_v$ change immediately and are proportional to the difference between $x_p^I$ and $x_v^I$ on each trial, so that they ramp down to zero in parallel with the perturbation. This contrasts with the gradual perceptual realignment changes observed in walking adaptation (Leech et al., 2018a; Vazquez et al., 2015) (also see our Control Experiments).

Additionally, the simulation of the mapping mechanism, $G^* = x_p^{per}(k) + \frac{1}{K}\left(p(k+1) - x_p(k+1)\right)$, failed to account for the motor results in this phase, exhibiting the same issues as the original PReMo (Fig. S7-8B, middle row resembles top row). This occurs because the overall motor output $x_p$, which includes both recalibration and mapping mechanisms, changes gradually according to the learning rate $K$. Consequently, changes in $G$ take many trials to be fully reflected in $x_p$.

Hence, we found complementary limitations where PReMo assumes perceptual realignment changes immediately while mapping adjustments develop gradually – but the opposite is true in our data. To address these limitations, we introduced an update equation for $\beta_p$ so that it changes gradually trial-by-trial according to the learning rate $K$. We then removed the learning rate from the update equation for $x_p$ so that it integrates two distinct types of changes: 1) the gradual changes in $x_p^{per}$ – driven by $\beta_p$ and representing the recalibration mechanism, and 2) the immediate changes in $G$ – representing the mapping mechanism. The final equations for the PM-ReMap model are reported in the main text. Note that setting $K = 0$ in the Ramp Down phase captures the special case of no unlearning or forgetting of recalibration. In this case, the model reduces to a single parameter $\eta_p$, representing the extent of perceptual realignment, and becomes mathematically equivalent to the recalibration + mapping model.

For the simulations, we set $W_p = \frac{\sigma_u^2}{\sigma_u^2+\sigma_p^2} = \frac{1}{3}$, $W_v = \frac{\sigma_u^2}{\sigma_u^2+\sigma_v^2} = \frac{1}{3}$, and $\eta_p = 0.56$ like before. We set $K = 0.01$ as this leads to the same recalibration learning rate as previous simulations (78% of the total recalibration-driven adaptation is accomplished in 3 minutes). We first simulated implicit adaptation ($G = 0$) with no perceptual shift for step length asymmetry ($\eta_v = 0$), and found that the PM-ReMap model accurately predicts perception of equal speeds halfway through the Ramp Down task (Fig. S7A, bottom row). When accounting for the mapping mechanism, the PM-ReMap model accurately predicts the Ramp Down motor results (Fig. S7B, bottom row). Implementation of a mapping mechanism is possible because of the separable range of ideal $G^*$ values in the first versus second halves of the Ramp Down, a characteristic that was lacking from PReMo (Fig. S8B, compare bottom row to top and middle rows).

We finally evaluated different values of $\eta_v$. For $\eta_v = 0$, the PM-ReMap model accurately predicts the relationship between perceptual and motor results – predicting that belt speeds feel equal when the motor aftereffects first emerge. In contrast, for $\eta_v > 0$ it inaccurately predicts perception of equal speeds earlier in the task (Fig. S8A, bottom row, the “belt speed feel equal” configuration is progressively earlier for larger $\eta_v$ values, depicted in lighter green and yellow). A $\eta_v$ value of zero signifies that proprioceptive realignment occurs for motor output but not for step length asymmetry, consistent with predictions from our previous framework (Rossi et al., 2021). The motor output is thought to be the signal generating sensory predictions and may therefore recalibrate in response to errors in this prediction. In contrast, step length asymmetry is thought to reflect a proprioceptive observation of the motor outcome, and purely proprioceptive signals are not thought to recalibrate in split-belt adaptation (Rossi et al., 2021; Vazquez et al., 2015).

**Supporting Text**

<u>Ramp Down comparison between Experiments 1 and 2</u>

We performed supplementary analyses to evaluate whether the step length asymmetry data in the Ramp Up & Down task of Experiment 2 is consistent with the recalibration + mapping hypothesis (Fig. S9-10).

Figure S9A-B show step length asymmetry time courses for Experiment 1 Ramp Down and Experiment 2 Ramp Up & Down. The magenta portion of the Ramp Up & Down task of Experiment 2 consists of the same speeds as the entire Ramp Down of Experiment 1 – i.e., speed differences ramping down from 1m/s to 0m/s. Despite the speeds being the same, we do not expect the step length asymmetry data to be the same in the two experiments. This is because participants in Experiment 2 are exposed to larger speed differences (1m/s to 1.5m/s) in the preceding, teal portion of the Ramp Up & Down. As these speed differences are larger than the adaptation speed difference (1m/s), additional learning is thought to occur.

We specifically evaluate whether the pattern of aftereffects differs between experiments. Participants in Experiment 1 walk symmetrically for speed differences ranging from 1m/s to 0.5m/s ("no aftereffect" range in Fig. S9A), and this absence of aftereffects is a key feature supporting the recalibration + mapping hypothesis. We therefore evaluate aftereffects in the same speed range for Experiment 2 ("no aftereffect in E1" range in Fig. S9B). In contrast to Experiment 1, participants in Experiment 2 appear to have positive step length asymmetry in this same speed range. Indeed, a statistical analysis confirmed that aftereffects were presents for 7 of these 11 speed configurations (see Table S8; we compared step length asymmetry for each speed configuration to zero using the same analysis as Experiment 1).

We propose that the presence of additional aftereffects in Experiment 2 is consistent with the additional learning occurring in the teal portion of the Ramp Up & Down. We performed two analyses to formally support this interpretation and confirm that the data is best explained by combination of recalibration and mapping mechanisms.

*Analysis 1: the aftereffect magnitude is correlated with the extent of additional learning*

In our first analysis, we formally assessed the relationship between the additional aftereffects and the additional learning observed in Experiment 2.

We define "additional aftereffect" as the mean step length asymmetry over strides taken at speed differences ranging from 1m/s to 0.5m/s (Fig. S9B, "no aftereffect in E1" range). Participants in

Experiment 1 exhibited no significant aftereffects in this range, so the “additional aftereffect” reflects aftereffects present in Experiment 2 but not Experiment 1, capturing the differences between experiments that we aim to study. A statistical analysis confirmed that there was a significant group-level additional aftereffect in Experiment 2 that was not present at the same speeds in Experiment 1 (mean SLA over speed differences in range 1m/s to 0.5m/s for Experiment 2 = 0.030 [0.009 0.051], for Experiment 1 = 0.008 [-0.014 0.029], group mean [CI]).

We define “additional learning” as the mean step length asymmetry over strides taken at speed differences ranging from 1m/s to 1.5m/s – corresponding to the teal portion of the Ramp Up & Down (Fig. S9B, “larger than adaptation” range). We consider the simplest scenario where any compensation for the larger speed difference occurs via additional learning (we discuss structure-based extrapolation and perform subgroup analyses below). Without additional learning, participants would exhibit pronounced negative step length asymmetry in this portion, as the belt speed difference is larger than during adaptation. Conversely, if they fully learned to account for the largest 1.5 m/s speed difference, they may also show aftereffects in the teal portion of the task as the speed difference ramps back down from 1.5 m/s to 1 m/s, resulting in positive step length asymmetry. Therefore, this measure is not expressed relative to zero but captures the relative extent of learning, with less negative or more positive values indicating greater learning.

Figure S9C shows the "additional aftereffect" versus "additional learning" measures for individual participants in Experiment 2. As expected, these measures appeared to be related: participants with larger additional aftereffects in the magenta portion of the task typically underwent more additional learning in the preceding teal portion. Pearson's correlation coefficient confirmed a significant correlation between the additional aftereffect and additional learning measures ($r=0.57$, $p=0.008$). This supports our interpretation that the different pattern of aftereffects in Experiment 2 versus Experiment 1 (for speed differences smaller than 1 m/s) may be due to the additional exposure to larger speed differences (>1m/s) present only in Experiment 2.

*The additional aftereffect - additional learning pattern is driven primarily by the memory-based subgroup*

The pattern of additional aftereffect and additional learning described above may differ between memory-based and structure-based subgroups of Experiment 2. The structure-based subgroup may compensate for the larger belt speed differences in the teal portion of the task using structure-based extrapolation of the stimulus-response mechanism, a process that differs from “actual” learning as it does not contribute to additional aftereffects in the magenta portion of the task. However, our "additional learning" measure does not distinguish between the processes, but captures the overall compensation achieved by actual learning as well as extrapolation. Therefore, participants in the structure-based subgroup may have larger additional learning without associated additional aftereffects. As such, we hypothesized that our group-level results for Analysis 1 were driven primarily by the memory-based subgroup.

To test this, we repeated Analysis 1 separately for the memory-based and structure-based subgroups. As expected, the analysis performed on the memory-based subgroup produced results consistent to that on the entire group: the additional aftereffect measure was significantly different from zero and significantly correlated with the additional learning measure for the memory-based subgroup (additional aftereffect = 0.028 [0.005 0.055], subgroup mean [CI]; subgroup correlation between additional aftereffect and additional learning: r=0.65, p=0.024). In contrast, results for the structure-based subgroup differed from those for the entire group: the additional aftereffect measure was not significantly different from zero nor correlated with the additional learning measure for the structure-based subgroup (additional aftereffect = 0.033 [-0.002 0.063], subgroup mean [CI]; subgroup correlation between additional aftereffect and additional learning: r=0.68, p=0.062). In sum, the subgroup analysis corroborates the finding that the additional aftereffects present in Experiment 2 but not Experiment 1 are related to the additional learning thought to occur in response to larger belt speed differences when the stimulus-response mapping is memory based.

*Analysis 2: flexible recalibration + mapping model fits the data better than recalibration only*

In Analysis 2, we used a modelling analysis to assess whether the step length asymmetry data during the magenta portion of the Ramp Up & Down task of Experiment 2 (1m/s to 0m/s speed differences) best aligns with the 1) recalibration + mapping or 2) recalibration only hypothesis.

We formulated a "flexible" version of the recalibration + mapping model that could account for the additional learning occurring in the teal portion of the Ramp Up & Down (>1m/s speed differences). This learning is complex because the perturbation exposure has mixed duration, size, and schedule: the long abrupt exposure to a large perturbation in adaptation is followed by the short gradual exposure to a small additional perturbation in the teal Ramp Up & Down. We used information provided by the speed match control experiments on how this additional exposure may affect the recalibration and mapping mechanisms, and modified the recalibration

+ mapping model of Experiment 1 accordingly. The step-by-step process and rationale underlying the development of this model are detailed below in section "*Development of the "flexible recalibration + mapping" model".* The final model accounts for additional but incomplete learning in each mechanism, as well as for the changing relationship between the mechanisms:

$$\text{Flexible Recalibration + Mapping:} \quad u(p) = \begin{cases} r + \alpha[p - \gamma r] & , \ \text{for } p \geq r \\ r & , \ \text{otherwise} \end{cases}$$

Like the original model, $r$ is a parameter capturing the amount of recalibration, $p$ is the perturbation, and $u$ is the motor output. We change the limits of the parameter $r$ to account for additional learning by the recalibration mechanism. There are two new parameters capturing additional incomplete learning by the mapping mechanism: $\alpha$ captures learning of the *perturbation magnitude*, and $\gamma$ captures learning of the *relationship between recalibration and mapping*.

When $\alpha = 1$ and $\gamma = 1$, the model matches the original recalibration + mapping, representing the case where learning is complete: recalibration compensation = $r$, mapping compensation = $p - r$, total compensation = $p$. Smaller values of $\alpha$ represent cases where the mapping mechanism has not fully learnt to compensate for new perturbation magnitude. It only compensates for a proportion $\alpha$ of the full $p - r$ it is supposed to counter, leading to non-zero sloped step length asymmetry over the range $p \geq r$ (which was symmetric in Experiment 1). Smaller values of $\gamma$ represent cases where the mapping mechanism has not fully learnt to account for changes in the recalibration mechanism (due to additional learning and to the changing relationship between recalibration and mapping that arises from mixed perturbation types, see below). Therefore, mapping operates as if the recalibration extent was smaller than its true value – specifically, a proportion $\gamma r$ of its true value $r$ – which may lead to overcompensation and aftereffects.

Using this model, we carried out an analysis equivalent to that of Experiment 1. We used the dual state model to represent the recalibration only hypothesis (see Methods in the main text). We fit the flexible recalibration + mapping and the recalibration only models to the magenta portion of the Ramp Up & Down (i.e., the last 63 strides, with belt speed differences ranging from 1m/s to 0m/s). We compared the goodness of fit between models using BIC.

We show the data and model fits in Figure S10. As expected, the flexible recalibration + mapping model can capture the sharp increase in step length asymmetry slope when the speed difference reaches 0.5m/s (dashed vertical line in Fig. S10B, left). In contrast, the recalibration only model cannot explain this feature and instead models step length asymmetry as approximately constant in slope (Fig. S10B, right). We formally evaluated the model fits by computing the difference in BIC between the recalibration only and flexible recalibration + mapping models (Fig. S10C). We found that the flexible recalibration + mapping model fitted the data better than the recalibration only model for all 20 participants, and the BIC difference was statistically significant at group level (mean [CI] = 23.123 [18.336 , 28.001]). These results confirm that, despite differing from Experiment 1, the step length asymmetry pattern in the Ramp Up & Down task of Experiment 2 best aligns with the recalibration + mapping hypothesis.

*Development of the "flexible recalibration + mapping" model*

We designed a "flexible" version of the recalibration + mapping model that was based on the model used for Experiment 1, but accounted for additional learning by the recalibration and mapping mechanisms. The original recalibration + mapping model (eq. 3 in the main text) can be rewritten as follows:

Original Recalibration + Mapping: $$u(p) = \begin{cases} r + [p - r] & , \quad \text{for } p \geq r \\ r & , \quad \text{otherwise} \end{cases}$$

Where $r$ is the amount of learning by recalibration, and $[p - r]$ is the amount of learning by mapping. We incorporated additional learning by recalibration by changing the upper bound for the parameter $r$ to equal the magnitude of the perturbation at the peak speed difference of 1.5m/s (mean over the 3 strides at this speed) - in contrast to the perturbation magnitude in adaptation as in the original model.

In the original model, we assumed that the total learning was complete (equal to $p$ in adaptation), so that the amount of learning by mapping is not specified by a parameter but rather by the difference between total learning minus recalibration ($[p - r]$ ). Therefore, incorporating additional learning by mapping is more complicated than recalibration. In this process, we aimed to account for three key features of the teal Ramp Up & Down that may affect the behavior of the mapping mechanism. First, the exposure to larger speed differences is brief (~1 minute), so that learning is likely incomplete. Second, the exposure is gradual (the belt speed difference gradually increases from 1 to 1.5m/s). Third, the additional speed difference is small (the magnitude of the increase is 0.5m/s). Importantly, these features differ from adaptation, where the exposure is long (15 minutes), abrupt, and large (1m/s belt speed difference).

**Supporting Text**

We used results from our speed match control experiments (where we manipulated duration, schedule and magnitude of the exposure) to inform us on how to incorporate these features into the model. To account for the brief exposure time, we examined the results from the Short Ascend and Short Descend control experiments. After a brief 3-minute exposure to a 1 m/s speed difference, learning by the mapping mechanism was partial, resulting in a non-zero sloped step length asymmetry in the 1 m/s to 0.5 m/s speed difference range (Note that there was still a visible sharp change in slope for smaller speed differences as expected by the mechanism + mapping hypothesis; see Figure 7B, black and yellow traces). To account for incomplete learning by the mapping mechanism due to exposure time, we introduced a parameter $\alpha$:

Recalibration + Mapping
w/ update for exposure time: $$u(p) = \begin{cases} r + \alpha[p-r] & , \quad \text{for } p \geq r \\ r & , \quad \text{otherwise} \end{cases}$$

Here, $[p-r]$ is the amount of "remaining" perturbation not accounted for by the recalibration mechanism. The parameter $\alpha$ captures the proportion of this remaining perturbation that is accounted for by mapping. The case of $\alpha = 1$ represent the scenario where learning is complete: the model is equivalent to that used for Experiment 1 and step length asymmetry in the range of speeds $r$ to $p$ (equal to the 1m/s to 0.5m/s speed difference range in Experiment 1) is zero. Step length asymmetry in this range becomes non-zero and sloped for smaller values of $\alpha$ ($0 < \alpha < 1$), matching the control experiments.

To account for the exposure magnitude and schedule, we examined results from the Small Gradual control experiment. After a gradual exposure to a small 0.4m/s belt speed difference, the recalibration mechanism contributes to ~80% of the total learning, while mapping contributes to only ~20% of the learning (Fig. 8A, green). This is in stark contrast to the equal proportions observed after an abrupt exposure to a large 1m/s belt speed difference, where recalibration and mapping mechanisms each contribute to ~50% of the total learning (as seen in the Medium Ascend and Descend control experiments, orange and dark blue in Fig. 8A, and in Experiment 1, "perceptual total" in Fig. 4B). Overall, the relationship between recalibration and mapping is not static in Experiment 2 because of the mixed schedule - abrupt adaptation to a 1m/s speed difference followed by additional gradual adaptation to the 1.5m/s speed difference in the Ramp Up & Down. In adaptation, mapping learns to counter ~50% of the perturbation, accounting for the recalibration mechanism that counters the remaining ~50%. In the teal Ramp Up & Down, mapping must update the relationship with the recalibration mechanism: it must learn to counter only ~20% of the perturbation, accounting for a recalibration mechanism that counters the remaining ~80%. However, learning is incomplete, so that the mapping mechanism may account for less of the recalibration mechanism. To account for this, we introduced a parameter $\gamma$:

$$\text{Flexible Recalibration + Mapping:} \quad u(p) = \begin{cases} r + \alpha[p - \gamma r] & , \quad \text{for } p \geq r \\ r & , \quad \text{otherwise} \end{cases}$$

The parameter $\gamma$ captures the proportion of the recalibration mechanism that mapping has learnt to account for. The case of $\gamma = 1$ represent the scenario where learning is complete: the model is equivalent to the recalibration + mapping w/ update for exposure time model defined above, mapping accounts for the "ideal" amount for recalibration, and step length asymmetry in the range of speeds $r$ to $p$ is zero or negative and sloped depending on $\alpha$. Smaller values of $\gamma$ ($0 < \gamma < 1$) capture scenarios where mapping has not learnt to account for the full recalibration amount: mapping operates as if the recalibration was smaller than its true magnitude, so that it may overcompensate and counter a larger proportion of the perturbation ($[p - \gamma r]$ versus $[p - r]$), leading to positive step length asymmetry aftereffects in the $r$ to $p$ speed range.

**Supporting Text**

### Clustering analysis

*Clustering results for Experiment 1*

The primary clustering analysis was designed to match that of Experiment 2 (see Supplementary Methods). For each participant, we evaluated the number of strides in the Ramp Down with step length asymmetry above their own baseline CI (2-min baseline block), indicating aftereffects (all CI reported in Table S10). Using the same density-based analysis as Experiment 2, we found no separate clusters in this measure (Fig. S12A). Nevertheless, the analysis detected two outlier participants (Fig. S12B). Because these participants showed aftereffects in *fewer* strides than the rest of the group, and walking without aftereffects for multiple speeds is evidence for mapping, we did not believe these outliers affected our main finding that adaptation involves mapping in addition to recalibration. We formally verified this by recomputing all statistical analyses of Experiment 1 without these outliers; this confirmed that none of the statistical results were affected by outliers (statistical results in Table S11).

As a further control, we repeated the clustering analysis using all the different measures evaluated in Experiment 1 (Fig. S13): BIC difference between the recalibration + mapping and recalibration only (dual state) models; "r" parameter; $compensation_{\text{motor total}}$; and PSE upper and lower bounds. We did not find evidence of separate clusters of participants using any of these measures.

*Clustering methodology*

We used the MATLAB "dbscan" algorithm (Ester et al., 1996) - a density-based clustering algorithm that clusters data points based on their relative Euclidean distance (here, the data points are the individual participants' measures described in the manuscript). The algorithm does not take number of clusters as an input, instead it takes the following two inputs: 1) Epsilon, i.e. the maximum distance between data points for them to be considered neighbors, 2) MinPts, i.e. the minimum number of data points in a neighborhood for the neighborhood to be labelled a cluster. A neighborhood is an ensample of data points where each data point is a neighbor of at least one other data point in the neighborhood (i.e. each data point has a distance < Epsilon to at least one other data point in the neighborhood). Data points in neighborhoods whose size is less than MinPts are labelled as outliers.

In order to ensure the clustering analysis would not depend on our choice of parameters, we adapted previously developed algorithms (Naik Gaonkar and Sawant, 2013; Rahmah and

Sitanggang, 2016) to automate the selection of Epsilon and MinPts. The original algorithm takes one parameter ‘k’ as an input, where ‘k’ can be an integer in the range of 1 to number of participants. To be fully unbiased, we here run the algorithm for all potential values of ‘k’ (ranging from 1 to 20), and develop a methodology to automatically select the best set of parameters out of the 20 ‘k’ iterations. We provide pseudocode for the full algorithm below – in the SI Appendix section “Pseudocode for the clustering analysis” – and explain here the general steps involved in the algorithm.

The first step is to obtain a set of epsilon-MinPts pairs to be tested. Potential epsilons are estimated based on the average distance of each participant to their closest ‘k’ neighbors. The algorithm computes the average distance measure and sorts it in ascending order across participants. It then detects locations where there is a sharp increase in average distance measure (specifically, where slope change – or concavity - is greater than 1% of the average slope). The values of the average distance measure at these locations are taken as potential epsilons to be evaluated. The potential epsilons are sorted by the average distance concavity (so that those obtained from locations with the sharpest change in slope are evaluated first). For each potential epsilon, the associated MinPts parameter is computed as the average number of neighbors that participants would have if that epsilon was used.

The second step is to select the most appropriate epsilon-MinPts pair for the current ‘k’ from the shortlisted options. First, epsilon-MinPts pairs are eliminated if MinPts is less than 2 (i.e. single participants could be detected as a standalone group) or more than 10 (i.e. clusters would be forced to have more than 10 people, hence it would be impossible for the algorithm to detect more than one group). The algorithm then runs dbscan for each remaining pair of epsilon-MinPts. The output is taken to be the first parameter pair that results in the smallest number of outliers.

The third step occurs after steps 1 and 2 are repeated for all ‘k’ values, and its goal is to select the overall best epsilon-MinPts pair across all ‘k’ iterations. First, it shortlists epsilon-MinPts pairs that resulted in the smallest number of outliers. Among these, it selects the epsilon-MinPts pair that led to the smallest number of clusters. This provided unique solutions on our data.

### Pseudocode

#### Pseudocode for the clustering analysis.

```
Notation (A x B) indicates matrix size. N = 20, K = 20 (i.e. the number of participants)

a.  Initialize BestEpsilon (1 x K), BestMinPts (1 x K), BestNumberOutliers (1 x K), BestNumberClusters (1 x K)

b.  Repeats steps 1-12 for k=1:K:
    1.  Dst (N x N) = distance of between each pair of participants:
        for n1=1:N
           for n2=1:N
                Dst (n1,n2) = abs (measure(n1) – measure(n2) )
    2.  AvgDst (N x 1) = mean distance to the first 'k' neighbors (for each participant):
        for n=1:N
           AvgDst(n) = mean ( Dst(n,1:k) )
    3.  SortAvgDst (N x 1) = AvgDst sorted in ascending order
    4.  SortAvgDstD (N-1 x 1) = first derivative of SortAvgDst
    5.  SortAvgDstDD (N-2 x 1) = second derivative of SortAvgDst
    6.  iEpsilon (M x 1) = indexes i where SortAvgDstDD (i) > 1/100 * mean(SortAvgDstD)
        (size M is automatic based on step 6)
    7.  Epsilon (M x 1) = SortAvgDst(iEpsilon+1)
    8.  SortEpsilon (M x 1) = Epsilon (j) where SortAvgDstDD (j) is sorted in descending order (i.e. sort Epsilon
        by descending SortAvgDstDD values)
    9.  MinPts (M x 1):
        for m=1:M
           initialize NumberNeighbors (N x 1)
           for n=1:N
                NumberNeighbors(n) = number of elements for which Dst (n, : ) ≤ SortEpsilon(m)
          MinPts(m) = mean (NumberNeighbors)
    10. Eliminate SortEpsilon (j) and MinPts (j) for which MinPts (j) < 2 or MinPts (j) > 10
    11. NumberOutliers (Q x 1): (Q is the size of SortEpsilon and MinPts after step 10)
        for q=1:Q
           clusters = dbscan (measure, SortEpsilon (q), MinPts(q) )
           NumberOutliers(q) = number of participants not assigned to any clusters
                (e.g. NumberOutliers(q) = number of elements for which clusters == -1,
                 with -1 = the label for outliers)
           NumberClusters(q) = number of different clusters
                (e.g. NumberCluster(q) = number of unique clusters different than -1,
                 with -1 = the label for outliers)
    12. BestEpsilon(k), BestMinPts(k), BestNumberOutliers(k), BestNumberClusters(k) = SortEpsilon(q),
        MinPts(q), NumberOutliers(q), NumberClusters(q), where q is the smallest value for which
        NumberOutliers(q) == min(NumberOutliers)

c.  OutlierBestEpsilon (J x 1), OutlierBestMinPts (J x 1), OutlierBestNumberClusters (J x 1) = BestEpsilon(j),
    BestMinPts(j), BestNumberClusters(j) for all j such that BestNumberOutliers(j) ==
    min(BestNumberOutliers) (size J is automatic based on step 12)

d.  OverallBestEspilon (1 x 1), OverallBestMinPts (1 x 1) = OutlierBestEpsilon(j), OutlierBestMinPts(j) for j
    such that OutlierBestNumberClusters(j) == min(OutlierBestNumberClusters)
    (note that in our dataset, we could choose any j in the last step and the resulting clustering is the same.
    This may not be true for different datasets)
```

### Supplementary Tables

**Table S1. Experiment 1, CI of step length asymmetry for each speed in the Ramp Up and Ramp Down tasks.** 95% bootstrapped confidence interval was corrected for multiple comparisons using false discovery rate (Benjamini and Yekutieli, 2005), Ramp Up $\boldsymbol{\alpha}_{\textbf{corrected}} = \frac{\textbf{6 significant comparisons}}{\textbf{7 total comparisons}} * \mathbf{0.05} = \mathbf{0.0429}$, Ramp Down $\boldsymbol{\alpha}_{\textbf{corrected}} = \frac{\textbf{12 significant comparisons}}{\textbf{21 total comparisons}} * \mathbf{0.05} = \mathbf{0.0286}$. Corrected CIs significantly different from zero are highlighted. Left speed was constant at 0.5m/s.

| | right speed (m/s) | mean | 95% CI | Corrected CI |
|---|---|---|---|---|
| **Baseline Ramp Up** | 0.35 | 0.203 | [0.177, 0.231] | **{0.175, 0.232}** |
| | 0.4 | 0.078 | [0.049, 0.105] | **{0.048, 0.106}** |
| | 0.45 | 0.006 | [-0.017, 0.028] | - |
| | 0.5 | -0.042 | [-0.064, -0.020] | **{-0.065, -0.019}** |
| | 0.55 | -0.088 | [-0.126, -0.050] | **{-0.127, -0.049}** |
| | 0.6 | -0.123 | [-0.149, -0.098] | **{-0.150, -0.097}** |
| | 0.65 | -0.144 | [-0.171, -0.119] | **{-0.172, -0.118}** |
| **Post-adaptation Ramp Down** | 1.5 | -0.015 | [-0.038, 0.009] | - |
| | 1.45 | -0.011 | [-0.035, 0.014] | - |
| | 1.4 | -0.01 | [-0.030, 0.010] | - |
| | 1.35 | 0.002 | [-0.025, 0.029] | - |
| | 1.3 | 0 | [-0.027, 0.031] | - |
| | 1.25 | 0.015 | [-0.013, 0.045] | - |
| | 1.2 | 0.018 | [-0.004, 0.042] | - |
| | 1.15 | 0.005 | [-0.019, 0.029] | - |
| | 1.1 | 0.024 | [0.000, 0.052] | {-0.002, 0.056} |
| | 1.05 | 0.023 | [0.002, 0.044] | {-0.001, 0.047} |
| | 1 | 0.026 | [-0.001, 0.052] | - |
| | 0.95 | 0.038 | [0.014, 0.059] | **{0.011, 0.062}** |
| | 0.9 | 0.056 | [0.026, 0.083] | **{0.023, 0.087}** |
| | 0.85 | 0.067 | [0.036, 0.097] | **{0.032, 0.100}** |
| | 0.8 | 0.083 | [0.056, 0.109] | **{0.053, 0.113}** |
| | 0.75 | 0.112 | [0.084, 0.138] | **{0.081, 0.142}** |
| | 0.7 | 0.126 | [0.091, 0.160] | **{0.087, 0.164}** |
| | 0.65 | 0.154 | [0.117, 0.192] | **{0.113, 0.196}** |
| | 0.6 | 0.174 | [0.143, 0.206] | **{0.139, 0.209}** |
| | 0.55 | 0.204 | [0.166, 0.243] | **{0.162, 0.248}** |
| | 0.5 | 0.213 | [0.172, 0.254] | **{0.168, 0.259}** |

### Supplementary Tables

**Table S2. Control experiments, CI of step length asymmetry in the first and last stride of the speed match task** (first post-adaptation task). 95% bootstrapped confidence interval was corrected for multiple comparisons using false discovery rate (Benjamini and Yekutieli, 2005), first stride $\boldsymbol{\alpha}_{\textbf{corrected}} = \frac{\textbf{5 significant comparisons}}{\textbf{7 total comparisons}} * \mathbf{0.05} = \mathbf{0.0357}$, last stride $\boldsymbol{\alpha}_{\textbf{corrected}} = \frac{\textbf{2 significant comparisons}}{\textbf{7 total comparisons}} * \mathbf{0.05} = \mathbf{0.0143}$. Corrected CIs significantly different from zero are highlighted.

| | Group | mean | 95% CI | Corrected CI |
|---|---|---|---|---|
| **First stride** | ShortAscend | 0.4572 | [0.3554, 0.5612] | **{0.3474, 0.5685}** |
| | MediumAscend | 0.433 | [0.2786, 0.5999] | **{0.2712, 0.6125}** |
| | LongAscend | 0.3887 | [0.1106, 0.6414] | **{0.0909, 0.6569}** |
| | ShortDescend | -0.1385 | [-0.1846, -0.1001] | **{-0.1880, -0.0972}** |
| | MediumDescend | -0.0165 | [-0.0691, 0.0393] | - |
| | SmallAbrupt | -0.0308 | [-0.0693, 0.0030] | - |
| | SmallGradual | -0.0648 | [-0.1210, -0.0063] | **{-0.1240, -0.0023}** |
| **Last stride** | ShortAscend | -0.014 | [-0.0291, 0.0007] | - |
| | MediumAscend | 0.0021 | [-0.0396, 0.0445] | - |
| | LongAscend | 0.0224 | [0.0003, 0.0471] | {-0.0045, 0.0542} |
| | ShortDescend | -0.0352 | [-0.0695, -0.0025] | {-0.0778, 0.0052} |
| | MediumDescend | 0.0431 | [-0.0004, 0.0899] | - |
| | SmallAbrupt | 0.0306 | [-0.0005, 0.0617] | - |
| | SmallGradual | -0.0041 | [-0.0323, 0.0246] | - |

### Supplementary Tables

**Table S3. Control experiments, within-group difference between $\boldsymbol{compensation}_{\text{motor total}}$ and $\boldsymbol{compensation}_{\text{perceptual}}$.** 95% bootstrapped confidence interval was corrected for multiple comparisons using false discovery rate (Benjamini and Yekutieli, 2005), $\boldsymbol{\alpha}_{\textbf{corrected}} = \frac{\textbf{7 significant comparisons}}{\textbf{7 total comparisons}} * \mathbf{0.05} = \mathbf{0.05}$. Significant differences based on corrected CI (Efron and Tibshirani, 1994) are highlighted.

| **Group** | **Mean** | **95% CI** | **Corrected CI** |
|---|---|---|---|
| Short Ascend | 47.450 | [36.082, 59.494] | **{36.082, 59.494}** |
| Medium Ascend | 48.295 | [38.795, 57.273] | **{38.795, 57.273}** |
| Long Ascend | 55.565 | [44.325, 66.551] | **{44.325, 66.551}** |
| Short Descend | 36.252 | [27.434, 44.096] | **{27.434, 44.096}** |
| Medium Descend | 56.012 | [42.462, 69.950] | **{42.462, 69.950}** |
| Small Abrupt | 36.224 | [17.254, 55.718] | **{17.254, 55.718}** |
| Small Gradual | 15.824 | [3.975, 27.015] | **{3.975, 27.015}** |

### Supplementary Tables

**Table S4. Control experiments, within-group comparison of $\boldsymbol{compensation}_{\textbf{motor total}}$ to 100%.** 95% bootstrapped confidence interval was corrected for multiple comparisons using false discovery rate (Benjamini and Yekutieli, 2005), $\alpha_{corrected} = \frac{5\ significant\ comparisons}{7\ total\ comparisons} * 0.05 = 0.0357$. Corrected CIs significantly different from 100% are highlighted.

| Group | Mean | 95% CI | Corrected CI |
|---|---|---|---|
| ShortAscend | 73.200 | [65.330, 81.111] | **{64.812, 81.563}** |
| MediumAscend | 90.045 | [86.579, 93.965] | **{86.336, 94.239}** |
| LongAscend | 98.815 | [93.457, 104.583] | - |
| ShortDescend | 70.752 | [63.513, 77.437] | **{63.013, 77.843}** |
| MediumDescend | 91.012 | [84.132, 98.274] | **{83.758, 98.773}** |
| SmallAbrupt | 98.349 | [86.146, 111.269] | - |
| SmallGradual | 73.074 | [61.812, 85.213] | **{61.141, 85.994}** |

### Supplementary Tables

**Table S5. Control experiments, between-group differences in $\boldsymbol{compensation}_{\text{motor total}}$.** 95% bootstrapped confidence interval was corrected for multiple comparisons using false discovery rate (Benjamini and Yekutieli, 2005), $\alpha_{corrected} = \frac{3\ significant\ comparisons}{6\ total\ comparisons} * 0.05 = 0.025$. Significant differences based on corrected CI (Efron and Tibshirani, 1994) are highlighted.

| Groups | Mean | 95% CI | Corrected CI |
|---|---|---|---|
| MediumAscend - ShortAscend | 16.845 | [7.981, 25.580] | **{7.029, 26.780}** |
| LongAscend - MediumAscend | 8.770 | [2.197, 15.529] | **{1.126, 16.418}** |
| ShortDescend - ShortAscend | -2.448 | [-13.194, 7.983] | - |
| MediumDescend - MediumAscend | 0.967 | [-6.939, 9.079] | - |
| SmallAbrupt - MediumDescend | 7.336 | [-6.881, 21.821] | - |
| SmallGradual - SmallAbrupt | -25.275 | [-42.137, -7.837] | **{-44.829, -5.784}** |

### Supplementary Tables

**Table S6. Control experiments, between-group differences in recalibration contribution to total output** ($compensation_{\mathrm{perceptual}}$ / $compensation_{\mathrm{motor\ total}}$). 95% bootstrapped confidence interval was corrected for multiple comparisons using false discovery rate (Benjamini and Yekutieli, 2005), $\alpha_{corrected} = \frac{1\ significant\ comparison}{6\ total\ comparisons} * 0.05 = 0.0083$. Significant differences based on corrected CI (Efron and Tibshirani, 1994) are highlighted.

| Groups | Mean | 95% CI | Corrected CI |
|---|---|---|---|
| Medium Ascend – Short Ascend | 9.277 | [-5.880, 24.373] | - |
| Long Ascend – Medium Ascend | -2.049 | [-17.108, 12.192] | - |
| Short Descend – Short Ascend | 12.343 | [-3.167, 28.054] | - |
| Medium Descend – Medium Ascend | -7.303 | [-24.239, 9.448] | - |
| Small Abrupt – Medium Descend | 26.546 | [7.235, 47.436] | **{0.488, 54.744}** |
| Small Gradual – Small Abrupt | 16.238 | [-5.608, 38.497] | - |

### Supplementary Tables

**Table S7. Control experiments, within-group difference between $\mathit{compensation}_{\text{motor recalibration}}$ and $\mathit{compensation}_{\text{perceptual}}$ across washout.** 95% bootstrapped confidence interval was corrected for multiple comparisons using false discovery rate (Benjamini and Yekutieli, 2005). $\boldsymbol{\alpha}_{\textbf{corrected}} = \frac{\textbf{3 significant comparisons}}{\textbf{6 total comparisons}} * \mathbf{0.05} = \mathbf{0.0250}$ for Short Ascend; $\boldsymbol{\alpha}_{\textbf{corrected}} = \frac{\textbf{5 significant comparisons}}{\textbf{6 total comparisons}} * \mathbf{0.05} = \mathbf{0.0417}$ for Medium and Long Ascend. Significant differences based on corrected CI (Efron and Tibshirani, 1994) are highlighted.

| | Time post-adaptation (min) | Mean | 95% CI | Corrected CI |
|---|---|---|---|---|
| **Short Ascend** | 0 | -0.074 | [-0.238, 0.076] | - |
| | 1 | -0.173 | [-0.280, -0.074] | **{-0.296, -0.062}** |
| | 2 | -0.113 | [-0.197, -0.032] | **{-0.210, -0.022}** |
| | 4 | -0.074 | [-0.117, -0.033] | **{-0.123, -0.028}** |
| | 8 | -0.045 | [-0.094, 0.004] | - |
| | 16 | -0.038 | [-0.127, 0.060] | - |
| **Medium Ascend** | 0 | -0.033 | [-0.249, 0.209] | - |
| | 1 | -0.183 | [-0.379, -0.001] | {-0.384, 0.005} |
| | 2 | -0.231 | [-0.381, -0.067] | **{-0.387, -0.061}** |
| | 4 | -0.111 | [-0.190, -0.029] | **{-0.193, -0.027}** |
| | 8 | -0.216 | [-0.344, -0.094] | **{-0.349, -0.089}** |
| | 16 | -0.163 | [-0.255, -0.059] | **{-0.258, -0.055}** |
| **Long Ascend** | 0 | 0.218 | [-0.018, 0.456] | - |
| | 1 | -0.209 | [-0.374, -0.051] | **{-0.380, -0.044}** |
| | 2 | -0.14 | [-0.260, -0.034] | **{-0.265, -0.031}** |
| | 4 | -0.11 | [-0.195, -0.024] | **{-0.198, -0.021}** |
| | 8 | -0.126 | [-0.239, -0.017] | **{-0.243, -0.013}** |
| | 16 | -0.113 | [-0.213, -0.027] | **{-0.218, -0.025}** |

### Supplementary Tables

**Table S8. Experiment 2, CI of step length asymmetry for each speed in the Ramp Up and magenta Ramp Up & Down tasks.** 95% bootstrapped confidence interval was corrected for multiple comparisons using false discovery rate (Benjamini and Yekutieli, 2005), Ramp Up $\boldsymbol{\alpha}_{\mathbf{corrected}} = \frac{\mathbf{6\ significant\ comparisons}}{\mathbf{7\ total\ comparisons}} * \mathbf{0.05} = \mathbf{0.0429}$, magenta Ramp Up & Down $\boldsymbol{\alpha}_{\mathbf{corrected}} = \frac{\mathbf{17\ significant\ comparisons}}{\mathbf{21\ total\ comparisons}} * \mathbf{0.05} = \mathbf{0.0405}$. Corrected CIs significantly different from zero are highlighted. Left speed was constant at 0.5m/s.

| | right speed (m/s) | mean | 95% CI | Corrected CI |
|---|---|---|---|---|
| **Baseline Ramp Up** | 0.35 | 0.197 | [0.145, 0.260] | **{0.143, 0.262}** |
| | 0.4 | 0.054 | [0.025, 0.081] | **{0.023, 0.082}** |
| | 0.45 | 0.015 | [-0.017, 0.048] | - |
| | 0.5 | -0.039 | [-0.066, -0.010] | **{-0.067, -0.009}** |
| | 0.55 | -0.068 | [-0.090, -0.045] | **{-0.090, -0.044}** |
| | 0.6 | -0.096 | [-0.118, -0.076] | **{-0.119, -0.075}** |
| | 0.65 | -0.109 | [-0.131, -0.089] | **{-0.132, -0.088}** |
| **Post-adaptation magenta Ramp Up & Down** | 1.5 | 0.008 | [-0.009, 0.026] | - |
| | 1.45 | 0 | [-0.016, 0.018] | - |
| | 1.4 | 0.018 | [-0.001, 0.040] | - |
| | 1.35 | 0.03 | [0.007, 0.056] | **{0.006, 0.057}** |
| | 1.3 | 0.028 | [0.003, 0.054] | **{0.002, 0.055}** |
| | 1.25 | 0.038 | [0.015, 0.062] | **{0.014, 0.064}** |
| | 1.2 | 0.041 | [0.019, 0.063] | **{0.018, 0.064}** |
| | 1.15 | 0.029 | [-0.001, 0.054] | - |
| | 1.1 | 0.043 | [0.017, 0.074] | **{0.016, 0.076}** |
| | 1.05 | 0.043 | [0.016, 0.069] | **{0.014, 0.071}** |
| | 1 | 0.051 | [0.019, 0.086] | **{0.018, 0.087}** |
| | 0.95 | 0.068 | [0.038, 0.101] | **{0.037, 0.102}** |
| | 0.9 | 0.072 | [0.048, 0.098] | **{0.047, 0.100}** |
| | 0.85 | 0.074 | [0.049, 0.099] | **{0.048, 0.101}** |
| | 0.8 | 0.083 | [0.058, 0.113] | **{0.057, 0.114}** |
| | 0.75 | 0.096 | [0.068, 0.126] | **{0.067, 0.127}** |
| | 0.7 | 0.115 | [0.085, 0.146] | **{0.084, 0.148}** |
| | 0.65 | 0.135 | [0.101, 0.173] | **{0.100, 0.176}** |
| | 0.6 | 0.188 | [0.134, 0.251] | **{0.132, 0.254}** |
| | 0.55 | 0.174 | [0.131, 0.222] | **{0.129, 0.224}** |
| | 0.5 | 0.233 | [0.163, 0.313] | **{0.160, 0.318}** |

### Supplementary Tables

**Table S9. Experiment 2, clustering analysis measures and results for individual participants.** Left column: 95% confidence interval of each participant's baseline (second tied-belt block). Middle column: number of strides in the first portion of the Ramp Up & Down (speeds larger than adaptation) with step length asymmetry below the participant's own baseline CI. Right column: clustering classification (cluster 1, memory; or cluster 2, structure).

| Participant | BL mean [95% CI] | # strides < CI | Cluster |
|---|---|---|---|
| P1 | 0.027 [0.016, 0.039] | 21 | Cluster 2 (structure) |
| P2 | 0.009 [0.004, 0.015] | 42 | Cluster 1 (memory) |
| P3 | -0.007 [-0.014, -0.000] | 55 | Cluster 1 (memory) |
| P4 | -0.017 [-0.025, -0.008] | 14 | Cluster 2 (structure) |
| P5 | 0.094 [0.082, 0.107] | 58 | Cluster 1 (memory) |
| P6 | 0.072 [0.060, 0.084] | 50 | Cluster 1 (memory) |
| P7 | 0.004 [-0.002, 0.011] | 21 | Cluster 2 (structure) |
| P8 | 0.024 [0.015, 0.032] | 51 | Cluster 1 (memory) |
| P9 | -0.045 [-0.053, -0.037] | 44 | Cluster 1 (memory) |
| P10 | -0.020 [-0.028, -0.013] | 3 | Cluster 2 (structure) |
| P11 | 0.002 [-0.007, 0.010] | 60 | Cluster 1 (memory) |
| P12 | -0.043 [-0.055, -0.032] | 16 | Cluster 2 (structure) |
| P13 | -0.027 [-0.037, -0.016] | 40 | Cluster 1 (memory) |
| P14 | 0.007 [-0.009, 0.024] | 58 | Cluster 1 (memory) |
| P15 | -0.055 [-0.068, -0.043] | 4 | Cluster 2 (structure) |
| P16 | -0.008 [-0.018, 0.003] | 40 | Cluster 1 (memory) |
| P17 | -0.068 [-0.081, -0.055] | 16 | Cluster 2 (structure) |
| P18 | -0.034 [-0.042, -0.026] | 11 | Cluster 2 (structure) |
| P19 | -0.009 [-0.017, -0.000] | 56 | Cluster 1 (memory) |
| P20 | 0.008 [-0.007, 0.022] | 38 | Cluster 1 (memory) |

### Supplementary Tables

**Table S10. Experiment 1, clustering analysis measures and results for individual participants.** Left column: 95% confidence interval of each participant's baseline (second tied-belt block). Middle column: number of strides in the ramp down with step length asymmetry above the participant's own baseline CI. Right column: clustering classification (cluster 1 or outlier).

| Participant | BL mean [95% CI] | # strides > CI | Cluster |
|---|---|---|---|
| P1 | 0.013 [0.005, 0.021] | 38 | Cluster 1 |
| *P2* | *-0.002 [-0.009, 0.006]* | *16* | *outlier* |
| *P3* | *0.106 [0.092, 0.121]* | *13* | *outlier* |
| P4 | 0.005 [-0.008, 0.018] | 53 | Cluster 1 |
| P5 | 0.019 [0.007, 0.032] | 47 | Cluster 1 |
| P6 | -0.040 [-0.058, -0.024] | 57 | Cluster 1 |
| P7 | -0.009 [-0.019, 0.002] | 58 | Cluster 1 |
| P8 | 0.027 [0.017, 0.037] | 25 | Cluster 1 |
| P9 | -0.019 [-0.030, -0.009] | 37 | Cluster 1 |
| P10 | -0.007 [-0.016, 0.003] | 48 | Cluster 1 |
| P11 | -0.038 [-0.045, -0.032] | 29 | Cluster 1 |
| P12 | -0.012 [-0.025, 0.000] | 60 | Cluster 1 |
| P13 | 0.022 [0.013, 0.030] | 40 | Cluster 1 |
| P14 | -0.046 [-0.054, -0.039] | 40 | Cluster 1 |
| P15 | -0.001 [-0.010, 0.008] | 33 | Cluster 1 |
| P16 | -0.066 [-0.073, -0.059] | 47 | Cluster 1 |
| P17 | 0.027 [0.016, 0.038] | 59 | Cluster 1 |
| P18 | 0.011 [-0.001, 0.024] | 38 | Cluster 1 |
| P19 | -0.060 [-0.077, -0.043] | 49 | Cluster 1 |
| P20 | 0.048 [0.039, 0.057] | 30 | Cluster 1 |

### Supplementary Tables

**Table S11. Experiment 1, replication of statistical analyses after removal of the 2 outliers detected with the primary clustering analysis.** Measures are reported the same way as the original analysis (rows 1-7 report mean [95% CI] {corrected CI}; rows 8-9 report correlation coefficient and p value). Significant results are highlighted. Removing the outliers did not affect the statistical significance of any of the tests. UB = upper bound, LB = lower bound.

| Measure | Statistical Results |
|---|---|
| "Recalibration Only" (dual state) − "Recalibration + Mapping" BIC | 8.435 [3.302, 13.896] **{3.302, 13.896}** |
| "Memory of Errors" − "Recalibration + Mapping" BIC | 11.522 [5.769, 17.791] **{5.769, 17.791}** |
| "Optimal Control" − "Recalibration + Mapping" BIC | 19.125 [13.956, 24.512] **{13.956, 24.512}** |
| $compensation_{\text{motor total}} - compensation_{\text{perceptual UB}}$ | 30.574 [23.788, 37.855] **{23.788, 37.855}** |
| $compensation_{\text{motor total}} - compensation_{\text{perceptual LB}}$ | 58.074 [49.016, 67.602] **{49.016, 67.602}** |
| $compensation_{\text{motor recalibration}} - compensation_{\text{perceptual UB}}$ | -7.606 [-14.425, -1.198] **{-14.425, -1.198}** |
| $compensation_{\text{motor recalibration}} - compensation_{\text{perceptual LB}}$ | 19.894 [9.259, 30.553] **{9.259, 30.553}** |
| Correlation between $compensation_{\text{motor recalibration}}$ & $compensation_{\text{perceptual UB}}$ | r=0.59, **p=0.010** |
| Correlation between $compensation_{\text{motor recalibration}}$ & $compensation_{\text{perceptual LB}}$ | r=0.30, p=0.226 |

### Supplementary Tables

**Table S12. Experiment 2, responses to questionnaire.** Raw responses of individual participants (center column) sorted by subgroup (left column), and classification of responses (three rightmost columns) used for Fig. 10.

| | | ***Response to:*** Did you deliberately change how you walked to account for how fast the belts were moving? If so, describe how. *Note: deliberately means that you thought about and decided to move that way* | **deliberate?** | **relevant?** | **accurate?** |
|---|---|---|---|---|---|
| **Memory-Based** | P2 | I noticed that I was making heavy steps on the right foot which caused shifting in my upper body so I tried to control that a little bit, but not much | Y | N | - |
| | P3 | Yes, I tried to adjust how fast I changed foot while walking, and tried to shift more weight onto my left side which was walking at a slower pace. | Y | Y | N |
| | P5 | Yes, I need to change how I walk because the belt moves in different speed | Y | N | - |
| | P6 | Yes, I changed my pace to stay upright and not fall down | Y | N | - |
| | P8 | No | N | - | - |
| | P9 | Yes, I started to change how I walked when I realized the right belt was moving faster, and I had to compensate on my left to stabilize my body. | Y | N | - |
| | P11 | No. | N | - | - |
| | P13 | Yes. I tried to get into a rhythm so that I had consistently long right strides and short left strides. | Y | N | - |
| | P14 | Yes, I tried to bow out my right leg(for the period of the time where the right belt was moving faster) so as to give me additional balance. I also tried to sync up my right foot moving up off the track at the same or close time as my left foot hit the track for the portion of the block where the right track was moving faster | Y | Y | N |
| | P20 | Yes, as soon as I felt the right belt's speed increase, I tried to spend as much time leaning on my left leg as possible and wait for a little before switching to my right foot (and then spending less time on my right foot) | Y | Y | Y |
| **Structure-Based** | P1 | Matched duration of standing on each foot by moving the right leg faster than the left leg and at a wider angle (range of motion). | Y | Y | N |
| | P4 | YES, I put less pressure on my right leg so I wouldn't lose my balance as the belts were moving at different rates. | Y | N | - |
| | P7 | Tried to even out the limp caused by the different leg speeds | Y | Y | N |
| | P10 | Yes I deliberately put more weight on my left foot (on the belt that was slower) to provide more stability for my right foot that was moving faster. | Y | N | - |
| | P12 | I felt like I had to adjust my speed to keep up with the way the belts were moving differently, I was thinking about how the right belt was moving faster but I didn't really feel like I had control over my own gait | N | - | - |
| | P15 | Yes, I adjusted as if I was limping. | Y | Y | N |
